# A genetic map of the response to DNA damage in human cells

**DOI:** 10.1101/845446

**Authors:** Michele Olivieri, Tiffany Cho, Alejandro Álvarez-Quilón, Kejiao Li, Matthew J. Schellenberg, Michal Zimmermann, Nicole Hustedt, Silvia Emma Rossi, Salomé Adam, Henrique Melo, Anne Margriet Heijink, Guillermo Sastre-Moreno, Nathalie Moatti, Rachel Szilard, Andrea McEwan, Alexanda K. Ling, Almudena Serrano-Benitez, Tajinder Ubhi, Irene Delgado-Sainz, Michael W. Ferguson, Grant W. Brown, Felipe Cortés-Ledesma, R. Scott Williams, Alberto Martin, Dongyi Xu, Daniel Durocher

**Author notes:** Ridgeline Therapeutics, Hochbergerstrasse 60C, CH-4057 Basel, Switzerland. Department of Biochemistry and Molecular Biology, Mayo Clinic, 200 First Street SW, Rochester, MN 55905, USA. Repare Therapeutics, 7210 Frederick-Banting, Suite 100, St-Laurent, QC, H4S 2A1, Canada. Topology and DNA breaks group, Spanish National Cancer Research Centre (CNIO), Madrid 28029, Spain.

## Abstract

The response to DNA damage is critical for cellular homeostasis, tumor suppression, immunity and gametogenesis. In order to provide an unbiased and global view of the DNA damage response in human cells, we undertook 28 CRISPR/Cas9 screens against 25 genotoxic agents in the retinal pigment epithelium-1 (RPE1) cell line. These screens identified 840 genes whose loss causes either sensitivity or resistance to DNA damaging agents. Mining this dataset, we uncovered that *ERCC6L2*, which is mutated in a bone-marrow failure syndrome, codes for a canonical non-homologous end-joining pathway factor; that the RNA polymerase II component ELOF1 modulates the response to transcription-blocking agents and that the cytotoxicity of the G-quadruplex ligand pyridostatin involves trapping topoisomerase II on DNA. This map of the DNA damage response provides a rich resource to study this fundamental cellular system and has implications for the development and use of genotoxic agents in cancer therapy.

## INTRODUCTION

The preservation of genetic information is a fundamental cellular process that represents a formidable challenge given that, in each cell and on a daily basis, the DNA polymer is the subject of a wide variety of chemical and physical alterations. These alterations, known collectively as DNA damage, have the potential to cause loss of genetic information, block transcription, halt DNA replication, impair chromosome segregation, generate mutations or produce chromosome rearrangements (Ciccia and Elledge, 2010; Jackson and Bartek, 2009; Lindahl and Barnes, 2000). These deleterious outcomes underlie many pathological conditions, including cancer and neurodegenerative diseases and are a likely cause of cellular and organismal aging (Hoeijmakers, 2009).

DNA damage can arise from many sources: from spontaneous chemical reactions such as base hydrolysis or deamination; from the interaction of DNA with reactive metabolites such as oxygen radicals, formaldehyde or S-adenosyl methionine (Lindahl and Barnes, 2000; Pontel et al., 2015; Tubbs and Nussenzweig, 2017); from the unscheduled or aberrant action of enzymes that act on DNA such as DNA topoisomerases, nucleases or cytidine deaminases; or through the action of environmental or human-made agents that include ultraviolet (UV) light or cancer chemotherapeutics such as the topoisomerase I poison camptothecin (Ciccia and Elledge, 2010; Tubbs and Nussenzweig, 2017). DNA replication is also liable to generate DNA damage through the incorporation of aberrant nucleotides such as ribonucleotides, when the replisome encounters obstacles such as strand interruptions, non-B DNA structures such as G-quadruplexes; or when the replisome collides with a transcribing RNA polymerase, a set of conditions known as DNA replication stress (Gomez-Gonzalez and Aguilera, 2019; Zeman and Cimprich, 2014). The DNA at stalled replisomes can also be processed to generate DNA double-strand breaks (DSBs), single-stranded (ss) DNA gaps or other structures that can be recognized as DNA damage, in an attempt to stimulate resumption of DNA synthesis or to overcome obstacles (Neelsen and Lopes, 2015; Quinet et al., 2017).

To counteract the deleterious outcomes of DNA damage, a complex and interconnected network of processes detect, signal and repair DNA lesions, which is often referred to as the DNA damage response (Jackson and Bartek, 2009; Zhou and Elledge, 2000). At the centre of this genome quality-control system are the various DNA repair pathways. These include base excision repair (BER), which deals with damaged bases and uracil; nucleotide excision repair (NER), which tackles helix-distorting lesions such as UV-induced pyrimidine dimers; mismatch repair (MMR), which removes misincorporated deoxyribonucleotides; single-strand break repair (SSBR), which re-seals single-strand breaks (SSBs) and the three DSB repair systems: non-homologous end-joining (NHEJ), homologous recombination (HR) and microhomology-mediated end-joining (MMEJ) (Caldecott, 2008; Gupta and Heinen, 2019; Hustedt and Durocher, 2016; Lindahl and Barnes, 2000; Marteijn et al., 2014; Sfeir and Symington, 2015). Some DNA lesions require the concerted action of multiple DNA repair pathways. One example is the repair of interstrand DNA crosslinks (ICLs), which involves the dual incision of the ICL directed by a subset of the Fanconi Anemia (FA) group of proteins followed by HR, the bypass of the DNA lesion by translesion DNA polymerases and final removal of the remaining adduct by the NER machinery (Kottemann and Smogorzewska, 2013).

DNA damage detection also elicits the activation of signal transduction cascades that are initiated by the ATM, ATR and DNA-PK protein kinases (Blackford and Jackson, 2017). DNA damage signaling coordinates DNA repair, modulates cell cycle progression, controls DNA replication initiation and regulates a wide range of biological processes (Blackford and Jackson, 2017; Jackson and Bartek, 2009). Furthermore, since the response to DNA damage occurs on chromatin, DNA repair takes advantage of, and is modulated by, chromatin remodeling and histone modification pathways (Papamichos-Chronakis and Peterson, 2013). The presence of DNA damage can also be communicated outside the cells harboring the lesion. As an example, DNA damage can lead to the activation of pattern recognition receptors, which can signal the presence of damaged cells to the immune systems for their elimination (Dhanwani et al., 2018). Therefore, the response to DNA damage is far-reaching and modulates multiple aspects of cell biology.

The cytotoxic, antiproliferative and immunostimulatory properties of DNA damage have been harnessed for the treatment of cancer for decades, and DNA damaging agents remain at the core of the anti-cancer armamentarium (O’Connor, 2015; Pilie et al., 2019). DNA repair processes are also selectively targeted in cancer therapy, as exemplified by the ability of poly (ADP-ribose) polymerase (PARP) inhibitors to selectively kill HR-deficient cells (O’Connor, 2015; Pilie et al., 2019). There are likely many additional opportunities to exploit the DNA damage response for the benefit of cancer patients, given that genome instability is a hallmark of cancer and there is intense interest in developing new modulators of the DNA damage response (Hanahan and Weinberg, 2011).

Despite the fact that genome maintenance mechanisms have been extensively studied, important DNA repair factors are still being uncovered. Recent examples include shieldin, which promotes NHEJ and antagonizes end-resection (Setiaputra and Durocher, 2019), and HMCES, a universally conserved suicide enzyme that protects abasic sites in single-stranded (ss) DNA (Mohni et al., 2019). We also know much less about how DNA damage intersects with cellular processes that are not directly involved in the modulation of DNA repair. Therefore, there is scope in applying hypothesis-free and unbiased methods to uncover new biological features of genome maintenance control in human cells.

The emergence of CRISPR-based genetic screens has enabled genome-scale analyses of gene-gene and gene-drug interactions in human cells (Hart et al., 2015; Shalem et al., 2014; Shalem et al., 2015; Wang et al., 2015). It is therefore possible to chart the human response to genotoxic stress in an unbiased and comprehensive manner. Indeed, recent work has illustrated the power of such screens in identifying new vulnerabilities to PARP and ATR inhibitors as well as to agents like temozolomide, a DNA alkylator (Hustedt et al., 2019a; MacLeod et al., 2019; Wang et al., 2018; Zimmermann et al., 2018). However, as these screens were carried out in isolation, they do not provide a global view of the human DNA damage response nor are they amenable to gene function inference by methods such as genetic interaction similarity profiling (Costanzo et al., 2019). We therefore aimed to map a genetic network of the response to DNA lesions and genotoxic agents, by undertaking 28 genome-scale CRISPR screens in an immortalized human diploid cell line (RPE-1 hTERT) that assessed the response to a wide variety of genotoxic agents that include multiple cancer therapeutics such as ionizing radiation, camptothecin, cisplatin and doxorubicin. The result is a rich dataset that uncovers new DNA repair factors, offers new insights into the mechanism-of-action of genotoxic drugs, and brings to light additional connections between genotoxic stress and extranuclear processes.

## RESULTS

We undertook a set of 25 CRISPR/Cas9 dropout screens in an hTERT-immortalized RPE-1 cell line clone that expresses Flag-tagged Cas9 and in which the gene encoding p53 is knocked out (Noordermeer et al., 2018). To this set, we added three other screens in the same cell line that have been previously presented (Hustedt et al., 2019a; Hustedt et al., 2019b; Noordermeer et al., 2018). In total, the 28 screens covered 25 genotoxic agents and the main features of the screens (library, genotoxin, concentration, etc.) are summarized in Table S1.

The screens were initially carried out using the TKOv2 sgRNA library but during the course of the study, we migrated to employing TKOv3, an all-in-one library with superior performance (Hart et al., 2017). The screens are schematized in Figure 1A and were carried out essentially as described previously (Hustedt et al., 2019b; Zimmermann et al., 2018). Briefly, cells were infected with the lentiviral sgRNA library at low multiplicity-of-infection (∼0.35) and were then divided into control and treated groups. The control group was left untreated for the duration of the experiment and the treated group was grown in the presence of a sublethal dose of drug for approximately 10 population doublings (Table S1). In the case of ultraviolet (UV) or ionizing radiation (IR) screens, we irradiated cells every 3 d, at an LD20 dose (Table S1). Gene-level depletion scores were computed using DrugZ, which is optimized for chemogenomic CRISPR screens (Colic et al., 2019).

**Figure 1.**
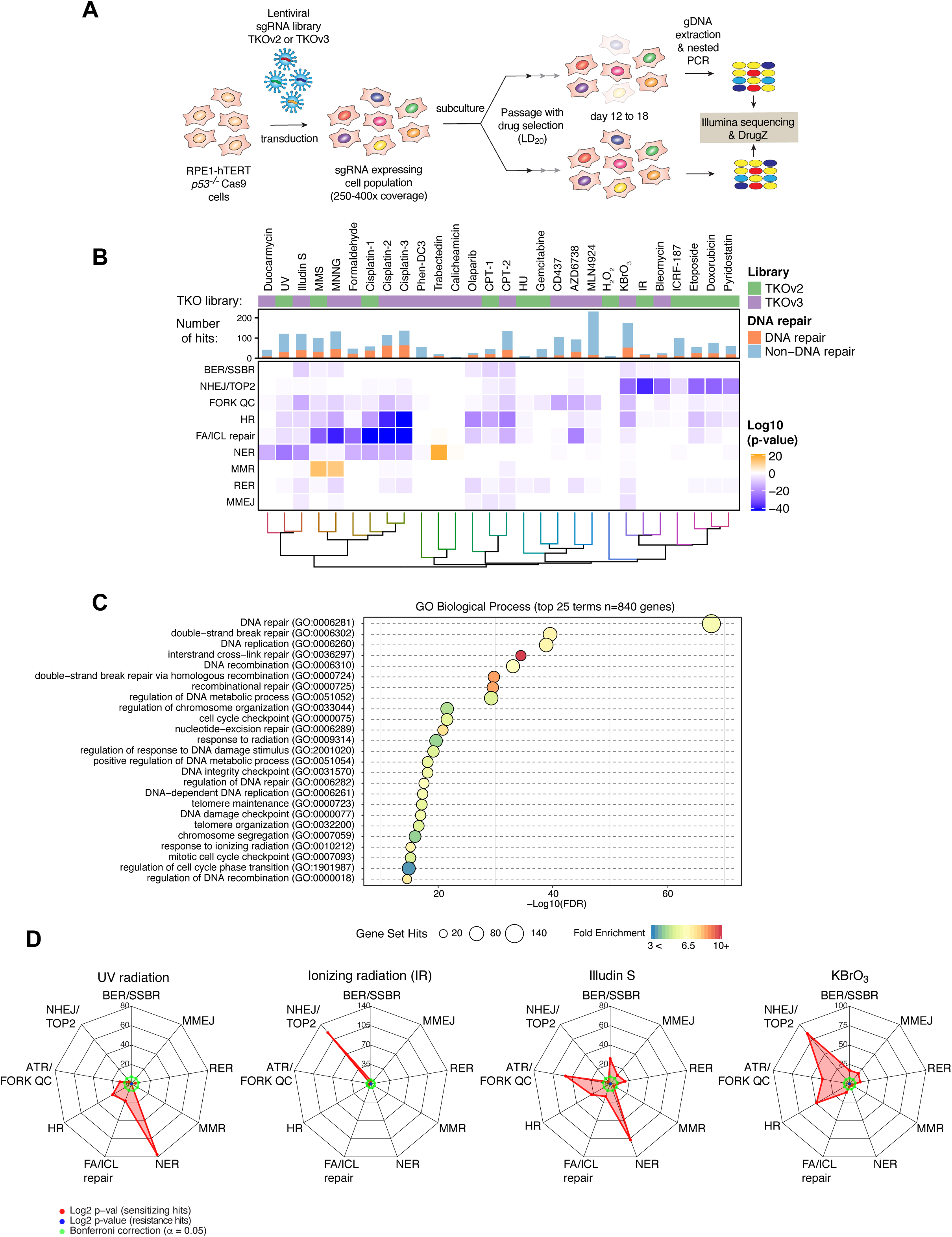
A chemogenomic view of the response to DNA damage. (A) Schematic of the dropout screens. (B) Heat map representation of the 28 CRISPR screens undertaken in RPE1 hTERT *p53^-/-^* Cas9 cells. The top panel indicates the library used. The histogram indicates the number of hits in each screen (determined as NormZ <-3 or >6, at FDR < 0.15) and the amount of DNA repair factor identified (curated list in Table S4). The lower heat map panel indicates the log_10_-transformed p values for enrichment of different DNA repair pathways (rows) calculated using a one-sided Fisher’s exact test. The color scale indicates fold-enrichment for resistance (orange) and sensitization (blue). Pathways are defined in the text except for ribonucleotide excision repair (RER). (C) Top 25 enriched GO terms, biological process, identified using g:Profiler (>10-fold enrichment; P < 0.05, with Benjamini-Hochberg FDR correction) among all 840 hits. Circle size indicates the number of genes from the common gene hit list included in each GO term, x-axis position indicates -log_10_-transformed FDR score and color indicates the fold enrichment compared with the whole-genome reference set. (D) Radar plots of genotoxic CRISPR screens performed with UV, IR, illudin S and KBrO_3_ depicting sensitization (red) or resistance (blue) for different DNA repair pathways (axes). Values indicate the log_2_-transformed p-values of the Fisher’s exact test score. Bonferroni thresholds are indicated in green. See also Figure S1.

The 25 genotoxic agents were chosen to cover most types of DNA damage, which range from DNA strand breaks (bleomycin, etoposide, IR, camptothecin), base alkylation (methyl methanesulfonate (MMS), and 1-methyl-3-nitro-1-nitrosoguanidine (MNNG)), inter- and intrastrand crosslinks (cisplatin, formaldehyde), oxidative DNA damage (H_2_O_2_, KBrO_3_), DNA replication stress (hydroxurea, CD437, gemcitabine), G-quadruplex stabilizing drugs (pyridostatin, Phen-DC3), transcription-interfering compounds (illudin S, trabectedin) and helix-distorting lesions (cisplatin, UV radiation). We also included profiles of DNA damage repair and signaling inhibitors such as olaparib (a PARP inhibitor), AZD6738 (an ATR inhibitor) and ICRF187 (a catalytic TOP2 inhibitor), as well as highly cytotoxic agents used in antibody-drug conjugates such as duocarmycin SA and calicheamicin (Table S1). The screens for cisplatin and camptothecin were carried out with both TKOv2 and TKOv3, which allowed us to ascertain the reproducibility of the screens across libraries and experimenters (Figure 1B and Table S1). The gene-level normalized Z-scores (NormZ) are presented in Table S2 and the read counts are presented in Datasets S1 and S2. Negative NormZ values represent genes whose mutation leads to their depletion from the cell population after genotoxin exposure, whereas positive NormZ scores represent genes whose mutation leads to a selective growth advantage in the presence of the drug. Each of the screens passed a series of quality-control tests described in the Methods section.

### Global view of the screens

We identified hits in the screen as follows: for genes whose mutation caused sensitization to the genotoxin, we selected NormZ values less than −3 with false discovery rates (FDR) lower than 15%. For genes whose mutation caused resistance to the genotoxic insult, we selected a NormZ value greater than 6. We employed this asymmetric set of thresholds since the screens were optimized to identify mutations that sensitize cells to genotoxins. These filters identified a total of 813 genes whose loss led to sensitization and 57 genes whose mutation led to resistance, in at least one screen, for a total of 840 unique genes. The median number of sensitizing mutations per screen was 56.5, ranging from 214 (for MLN4924) to only 1 for calicheamicin (Figure 1B and Table S2). In the latter case, this low number of hits was likely due to difficulties in maintaining an LD20 concentration of the drug over the duration of the screen.

Functional term enrichment analyses using g:Profiler (Raudvere et al., 2019) showed that the 840 genes are highly enriched in DNA repair-associated terms in all categories assessed, which included gene ontology (GO), KEGG, Reactome and WikiPathways (Table S3). For example, the top terms for the GO categories molecular function and biological process were “damaged DNA binding” (GO:0003684) and “DNA repair” (GO:0006281) respectively (Figures 1C, S1A and Table S3). The DNA repair associated terms were varied and covered most DNA repair processes ranging from DNA double-strand break repair to mismatch repair. Highly enriched terms not directly associated with DNA repair included DNA replication (GO:0006260), regulation of chromosome organization (GO:0033044) and covalent chromatin modification (GO:0016569) pointing to other processes that are known to influence the response to DNA damage. While the hits from the screens were highly enriched for DNA repair-associated terms, DNA repair terms only covered 336 of the 840 genes identified, suggesting that the screens identified genes not previously associated with the response to DNA damage or that may be associated with pathways that indirectly influence the response to genotoxic agents.

To more specifically help us derive insights into the DNA repair mechanisms involved in the response to the agents screened, we curated a set of 197 bona fide DNA damage repair and signaling factors grouped according to their known function (Table S4). This list did not include genes that were not represented in our sgRNA library or that were only weakly linked to DNA repair. We next determined whether these groups were enriched in each of the screens using a Fischer exact test with Bonferroni correction. To facilitate visualization, we represented the data as heatmaps (Figure 1B) or radar plots (Figures 1D and S1B). For example, NER was important for survival to UV radiation, as expected, with additional contributions of FA/ICL and HR pathway genes (Figure 1B,D). Similarly, genes promoting survival to IR were highly enriched in NHEJ factor-coding genes, in line with previous results indicating that NHEJ is a major contributor to the cellular resistance to IR (Figure 1B,D). Some less characterized agents, like Illudin S, showed that multiple pathways, such as HR and DNA replication fork quality control (QC) in addition to NER, are involved in mediating cellular resistance to this agent (Jaspers et al., 2002) (Figure 1B,D). A few surprises arose from this analysis. One example was the DNA repair sensitization profile of KBrO_3_, an oxidizing agent and suspected carcinogen used as food additive (IARC, 1999). KBrO_3_ is widely thought to oxidize DNA to produce 8-oxoguanine but whether this lesion is the root cause of bromate genotoxicity is unknown (Ballmaier and Epe, 2006). We found that KBrO_3_ cytotoxicity was greatly enhanced by the loss of genes acting in NHEJ, MMEJ and HR, in addition to the expected sensitization by loss of BER/SSBR factors (Figure 1B,D). These observations strongly suggest that DSBs are a relevant genotoxic lesion of KBrO_3_.

### Pyridostatin and CX5461 cytotoxicity involves TOP2 trapping

Another unsuspected insight into genotoxic drug mechanism of action concerns pyridostatin, a compound that avidly binds and stabilizes G-quadruplex sequences in vitro and in cells (Muller et al., 2010; Rodriguez et al., 2012). We profiled pyridostatin along with another G-quadruplex binder, Phen-DC3 (De Cian et al., 2007). Both compounds had very different profiles, with pyridostatin displaying clear sensitization following loss of NHEJ (Figure 1B and Figure S1B). Furthermore, when we performed hierarchical clustering of the compounds based on Pearson correlation coefficients (PCC) of either DNA repair genes (Figure 1B) or gene hits (Figure S2A), pyridostatin not only clustered with other clastogenic agents but also clustered closely to etoposide and doxorubicin, two agents that cause DSBs via the poisoning of topoisomerase II (TOP2) (Delgado et al., 2018). In fact, TDP2, NBS1 (NBN) and ZATT (ZNF451) were among the top 3 hits in the pyridostatin screen and all three proteins are involved in the removal of trapped TOP2 (Aparicio et al., 2016; Schellenberg et al., 2017) (Figure 2A). These observations hinted that the mechanism of pyridostatin cytotoxicity involves the poisoning of TOP2.

**Figure 2.**
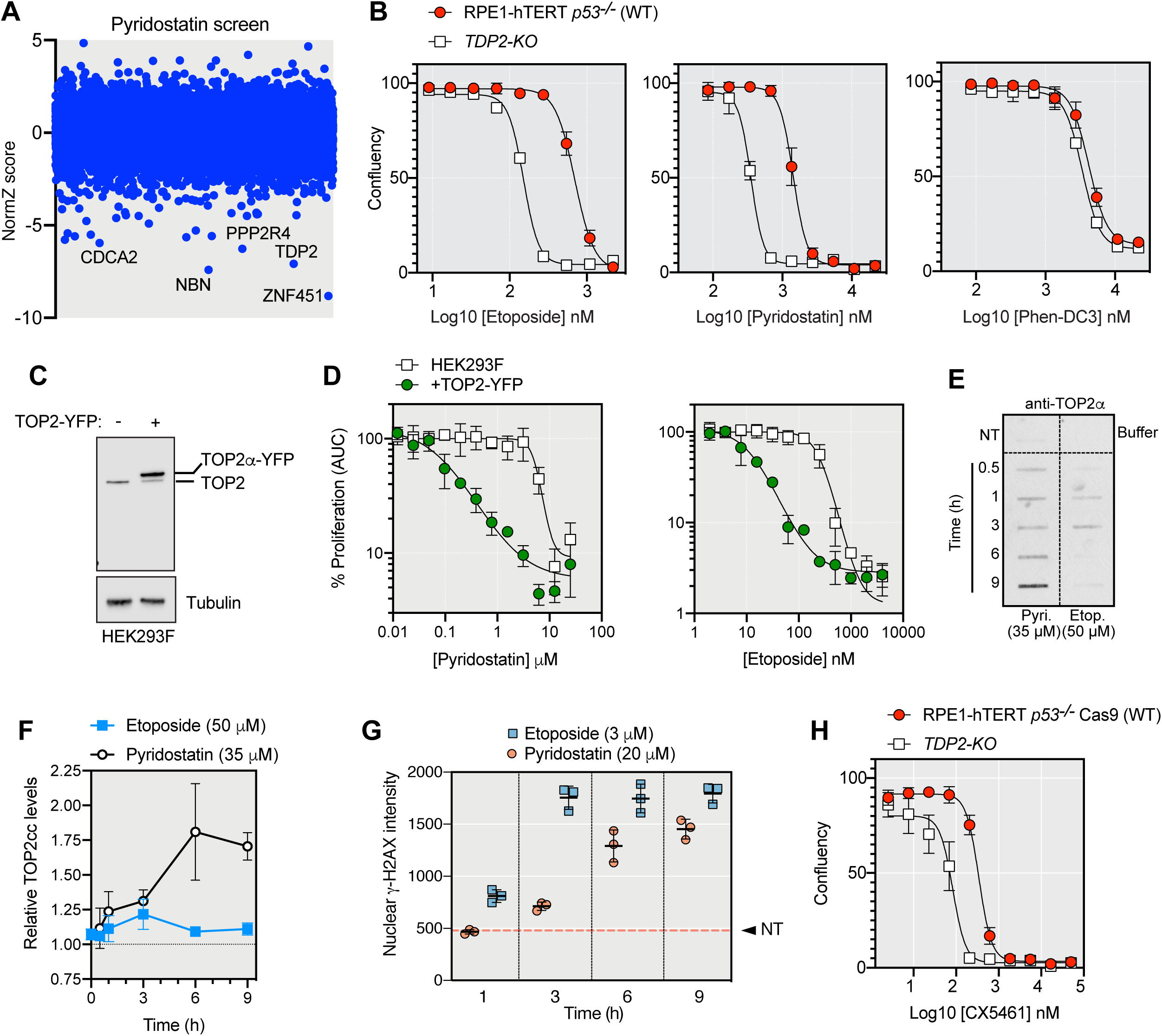
Pyridostatin and CX5461 cytotoxicity involves TOP2 trapping. (A) CRISPR dropout screen results for RPE1 cells exposed to pyridostatin. The 5 genes with the lowest NormZ values are labeled. (B) Drug-response assays with the indicated compounds in RPE1 and isogenic *TDP2^-/-^* cells using confluency as a readout 6 d post-treatment. Data presented as mean ± SD.; *N*=3. (C) Immunoblotting of TOP2 in HEK293F cells expressing TOP2-YFP or a control vector. Tubulin detection was used as loading control. (D) Drug-response assays with pyridostatin and etoposide in parental HEK293F cells and those expressing TOP2-YFP. Data presented as mean ± SEM; *N*=3. (E) RADAR assay for the detection of TOP2cc in RPE1 cells treated with etoposide (Etop.) and pyridostatin (Pyri.). The condition in upper left box represents samples that were untreated (NT). The upper right box represents buffer alone. (F) Quantitation of the RADAR assay normalized to untreated cells. Data presented as mean ± SD; *N*=3. (G) Average *γ*H2AX nuclear intensity in RPE1 cells treated with etoposide or pyridostatin determined by image segmentation. The dashed line represents the value for untreated (NT) cells. Data presented as mean ± SD; *N*=3. See also Figure S2.

To test this possibility, we confirmed that loss of TDP2 hypersensitizes cells to pyridostatin but not to Phen-DC3 (Figure 2B). The extent of sensitization by TDP2 loss was similar between pyridostatin and etoposide, a bona fide TOP2 poison (Figure 2B). Furthermore, overexpression of TOP2*α* fused to YFP in HEK293F cells, also led to sensitization to both etoposide and pyridostatin (Figures 2CD) consistent with the idea that cytotoxicity of pyridostatin involves TOP2. Assessment of TOP2 cleavage complexes (TOP2cc) using the RADAR assay (Kiianitsa and Maizels, 2014) showed that pyridostatin is a potent inducer of TOP2cc (Figure 2E). The kinetics of TOP2cc formation was strikingly different between etoposide and pyridostatin, with the etoposide-induced TOP2cc being barely above the detection threshold and displaying a biphasic accumulation characterized by a peak of TOP2cc 3 h post treatment (Figure 2EF). In contrast, pyridostatin caused TOP2cc to accumulate over the duration of the experiment (9 h), reaching levels of TOP2cc that were higher than those caused by etoposide (Figure 2EF). However, pyridostatin was much less potent at evoking *γ*-H2AX than etoposide, with pyridostatin-induced *γ*-H2AX formation being delayed and blunted compared to the response caused by etoposide, which was used at lower concentrations (Figure 2G and Figure S2B). These data suggest that pyridostatin-induced TOP2cc may be converted less efficiently into DSBs than those induced by etoposide, pointing to different mechanisms of TOP2 poisoning. Since *γ*-H2AX formation in pyridostatin-treated cells tends to occur in regions with propensity to form G-quadruplex structures (Rodriguez et al., 2012), one attractive possibility is that pyridostatin promotes TOP2 trapping following G-quadruplex stabilization.

As a means to see if these results could extend to other G-quadruplex ligands, we considered CX5461, which was originally identified as an RNA polymerase I inhibitor but was recently shown to be a G4 ligand (Xu et al., 2017). Like pyridostatin, CX5461 shows activity against HR-deficient cells and tumors (Xu et al., 2017; Zimmer et al., 2016), suggesting a common mechanism of action. We found that TDP2 loss hypersenstized cells to CX5461 (Figure 2H), consistent with the possibility that the cytotoxicity of CX5461 involves the trapping of TOP2.

### Gene-level view of the screens

To visualize the activity of genes across screens, we generated fingerprint plots that give a digital view of the gene activity across various screens (Figure 3A). As examples, we observed that the transcription-coupled (TC)-NER factor ERCC8 (also known as CSA), promotes cellular resistance to UV, illudin S and cisplatin, as expected (Figure 3A). In another example, the core NHEJ factors XRCC4 and LIG4 promoted survival against DSB-inducing agents such as IR, bleomycin and etoposide (Figure 3A). Some hits had exquisitely restricted profiles. For example, ATAD5, also known as ELG1, promotes cellular resistance exclusively to the alkylating agents MMS and MNNG (Figure 3A). ATAD5 forms an alternative RFC complex that may be involved in removing DNA-bound PCNA (Kanellis et al., 2003; Kang et al., 2019). Intriguingly, DSCC1 and CHTF8, which are components of another alternative RFC complex, RFC-CTF18, promote resistance to a wide variety of genotoxins and represent the group of genes that scored as hits in the highest number of screens (Figure 3A and Figure S3AB).

**Figure 3.**
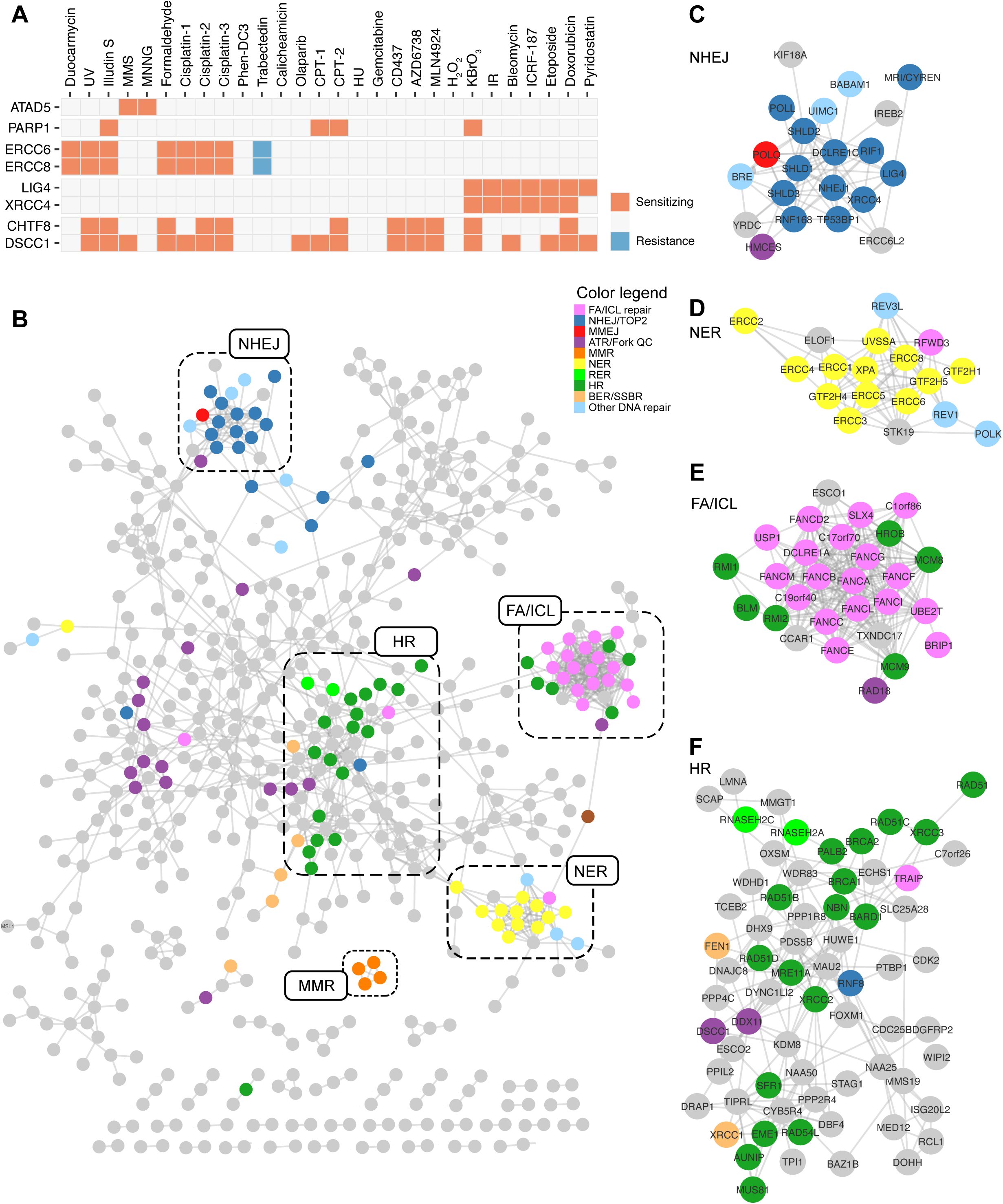
The DNA damage response network. (A) Fingerprint plot of highlighted genes across the 28 screens (columns). The boxes are labeled according to whether mutations in these genes leads to sensitization (orange) or resistance (blue). (B) Network of genes displaying at least one connection with another gene at a PCC value >0.725 visualized in (cytoscape v3.7.1) using a perfused force directed layout. Nodes are colored according to a DNA repair pathway curation (Table S4). Highlighted are the clusters enriched in DNA repair genes. The same network with gene names can be found in Figure S3C. (C-F) Details of the clusters enriched in NHEJ (C), NER (D), FA/ICL (E) or HR (F) pathway coding genes. See also Figures S3 and S4.

The similarity in profile of genes having related function (e.g. *XRCC4* and *LIG4* or *ERCC6* and *ERCC8*) prompted us to undertake similarity profiling based on PCC. This allowed us to build a network where genes are nodes and their edges are constrained by their PCC values (Figure 3B and Table S5; see Figure S3C for a network with gene names included). In order to better visualize structure within this network, we employed a high PCC threshold of 0.725 in Figure 3B. The resulting 568-gene network is composed of a highly connected core subnetwork with clearly defined submodules, many of which are highly enriched in DNA repair pathways such as NHEJ, NER and ICL repair that are connected directly or indirectly to a central portion enriched in HR factors (Figure 3C-F). The DNA repair-associated genes composed only 42% of the total network (240/568), suggesting that uncharacterized DNA repair factors are included in this network, and that other pathways are represented. As one example, we observed that Hippo pathway components such as NF2, FRYL, AMOTL2, LATS2 and TAOK1 clustered together in the network and their loss sensitized cells to multiple DNA damaging agents (Figures S4). This observation is consistent with the Hippo pathway acting as a major modulator of chemosensitivity to cancer therapeutics (Nguyen and Yi, 2019) and it suggests that additional Hippo signaling modulators could be found connected to these genes. Candidates for such proteins include KIRREL and FAM49B, which are connected to multiple known Hippo signaling molecules in the network (Figure S4). Therefore, we expect that exploration of this network will reveal insights about a wide range of pathways that impinge on survival to genotoxic stress including, but not limited to, DNA repair.

### The bone marrow failure syndrome gene *ERCC6L2* encodes an NHEJ factor

We next mined the DNA damage network to uncover uncharacterized or understudied DNA repair factors. One clear example resided in a subcluster enriched in NHEJ factor-coding genes (Figure 3C). This subcluster contained many core NHEJ factors (XRCC4, LIG4, NHEJ1), 53BP1-pathway modulators such as 53BP1, RIF1 and the shieldin components SHLD1-3 as well as the NHEJ regulator MRI/CYREN (Arnoult et al., 2017; Hung et al., 2018; Setiaputra and Durocher, 2019). This subcluster also contained ERCC6L2, a SWI/SNF-like ATPase remotely related to ERCC6 (CSB) and ERCC6L (PICH) (Figure 4A). *ERCC6L2* was of particular interest since its mutation causes an inherited bone marrow failure (iBMF) syndrome, myelodysplastic syndrome and leukemia (Douglas et al., 2019; Jarviaho et al., 2018; Shabanova et al., 2018; Tummala et al., 2014; Zhang et al., 2016). ERCC6L2 deficiency is also associated with additional clinical phenotypes such as microcephaly, ataxia and other developmental delays that are often associated with NHEJ deficiency (Nijnik et al., 2007; Shabanova et al., 2018; Woodbine et al., 2014). While ERCC6L2 has previously been associated with genome instability, there is currently debate as to which DNA repair pathway it participates in, with most recent results pointing to ERCC6L2 participating in the resolution of R-loops (Tummala et al., 2018).

**Figure 4.**
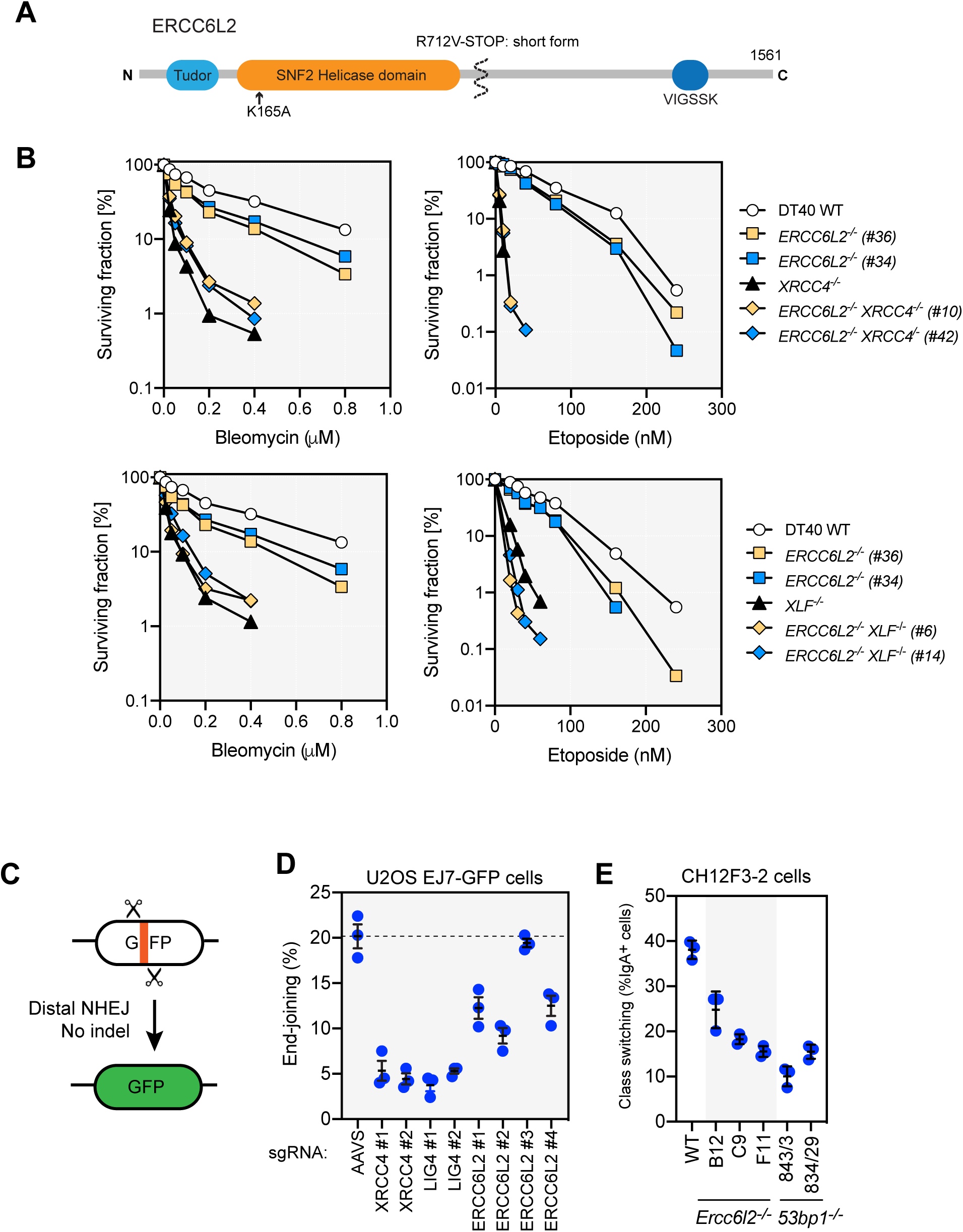
ERCC6L2 promotes canonical NHEJ. (A) Schematic overview of ERCC6L2. ERCC6L2 also possess a short isoform of 712 residues produced by alternative splicing. (B) Cell proliferation assays of DT40 cells of the indicated genotypes treated with either etoposide or bleomycin for a 3 d period. Data is presented as the mean of a technical triplicate. An independent experiment is shown in Figure S5C. (C) Schematic of the EJ7-GFP NHEJ assay. (D) End-joining frequency of U2OS-EJ7 cells transduced with sgRNA-expressing viruses targeting the indicated genes and subsequently transfected with a vector expressing an sgRNA targeting the *EGFP* stuffer region in the reporter transgene. Dashed line represents the mean of the end-joining frequency of the *AAVS1*-targeted condition. Error bars represent the mean ± SD; *N*=3. TIDE analysis is shown in Table S6. Note that sgERCC6L2-3 has low indel efficiency. (E) Class switching recombination levels (% IgA^+^ cells) in CH12F3-2 cells of the indicated genotypes. Error bars represent the mean ± SD; *N*=3. See also Figure S5.

To assess the role of ERCC6L2 in DNA repair, we first generated independent knockout clones in the chicken DT40 cell line (Figure S5AB). The *ERCC6L2* knockouts (*ERCCL62*^-/-^) were constructed in wild-type, *XLF^-/-^* and *XRCC4^-/-^* cell backgrounds to assess a possible genetic relationship between *ERCCL2* and core NHEJ factors (Xing et al., 2015). As expected from the screens, loss of ERCC6L2 led to sensitivity to etoposide and bleomycin although the extent of sensitization was milder than that caused by null mutations in *XLF* or *XRCC4* (Figure 4B). The *ERCC6L2*^-/-^ *XRCC4*^-/-^ double-mutant lines consistently showed similar sensitivity to bleomycin or etoposide as *XRCC4^-/-^* cells, indicating that *ERCC6L2* acts in the same genetic pathway as *XRCC4* (Figure 4B and Figure S5C). The situation was similar with XLF deficiency, where the *ERCC6L2*^-/-^ *XLF^-/-^* double-mutant cells showed a similar sensitivity profile as *XLF^-/-^* cells in response to bleomycin, but the double-mutant cells were slightly more sensitive to etoposide than *XLF^-/-^* cells (Figure 4B and S5C). These data indicate that ERCC6L2 acts in the canonical XRCC4-dependent NHEJ pathway but can provide support to NHEJ independently of XLF under certain conditions such as TOP2-driven DSBs.

In support for a role for ERCC6L2 in NHEJ, we employed the recently described EJ7-GFP reporter (Bhargava et al., 2018), which is unique among the many NHEJ assays by being highly dependent on canonical NHEJ factors, to assess the role of ERCC6L2 in this pathway in the human U2OS cell line. The assay monitors the end-joining of two distal Cas9-induced DSBs without indel formation (Figure 4C). We observed that the loss of ERCC6L2 following introduction of effective sgRNAs against its gene, led to a marked reduction in end-joining, albeit not as much as that caused by XRCC4 or LIG4 depletion (Figure 4D and Tables S6). These results are consistent with a role for ERCC6L2 in promoting canonical NHEJ.

To further assess the role of ERCC6L2 in physiological NHEJ, we examined the impact of its loss for immunoglobulin class switching (Methot and Di Noia, 2017) in mouse CH12F3-2 cells (Nakamura et al., 1996). We generated three independent *Ercc6l2^-/-^* clones and monitored switching from IgM to IgA by flow cytometry. All three *Ercc6l2* knockout clones showed reduced numbers of IgA^+^ cells following induction of switching, with the magnitude of the defect approaching that of *53bp1*^-/-^ cells, which are highly impaired in class switching (Figure 4E). The defect was not due to reduced expression of germline transcripts (*glt*) at the *Ig* locus, or AID, which are both necessary for DSB induction, hinting at a direct role in the repair of DNA breaks (Figure S5DE). Collectively, these data suggest that ERCC6L2 is a bona fide NHEJ factor and indicates that the root cause of the bone marrow failure in individuals with mutated *ERCC6L2* might be a failure to appropriately repair DSBs.

### ELOF1 and STK19 modulate TC-NER

The NER cluster in the PCC network contains primarily TC-NER factors, such as ERCC8/CSB, UVSSA and various TFIIH components (ERCC2, GTF2H1, GTF2H4 and GTF2H5). Consistent with this cluster reflecting primarily TC-NER, the global-genome (GG) NER factor XPC was not connected to this group of genes even though it was a strong hit in the UV screen (Figures 3D and Figure 5A). We noted two genes within this cluster that were not previously associated with TC-NER. The first is ELOF1, an RNA polymerase II-associated protein that is conserved across eukaryotes and present in archaea (Daniels et al., 2009). The ELOF1 homolog in budding yeast, Elf1, plays a role in transcriptional elongation and recent structural studies indicate that Elf1 collaborates with Spt5/6 (DSIF in human) to promote transcriptional elongation past the chromatin barrier (Ehara et al., 2019; Ehara et al., 2017; Prather et al., 2005). In the screens, loss of ELOF1 results in sensitivity to UV, illudin S, cisplatin (2 out of 3) as well as resistance to trabectedin (Figure 5A). Indeed, cells require the presence of TC-NER factors to survive the transcription-blocking agent illudin S; conversely, for the chemotherapeutic drug trabectedin, TC-NER factors are necessary for its cytotoxic effects (Takebayashi et al., 2001). This drug response profile of ELOF1 closely matched that of other TC-NER factors although the depletion scores for *ELOF1* were generally lower than canonical TC-NER factor-coding genes (Table S2).

**Figure 5.**
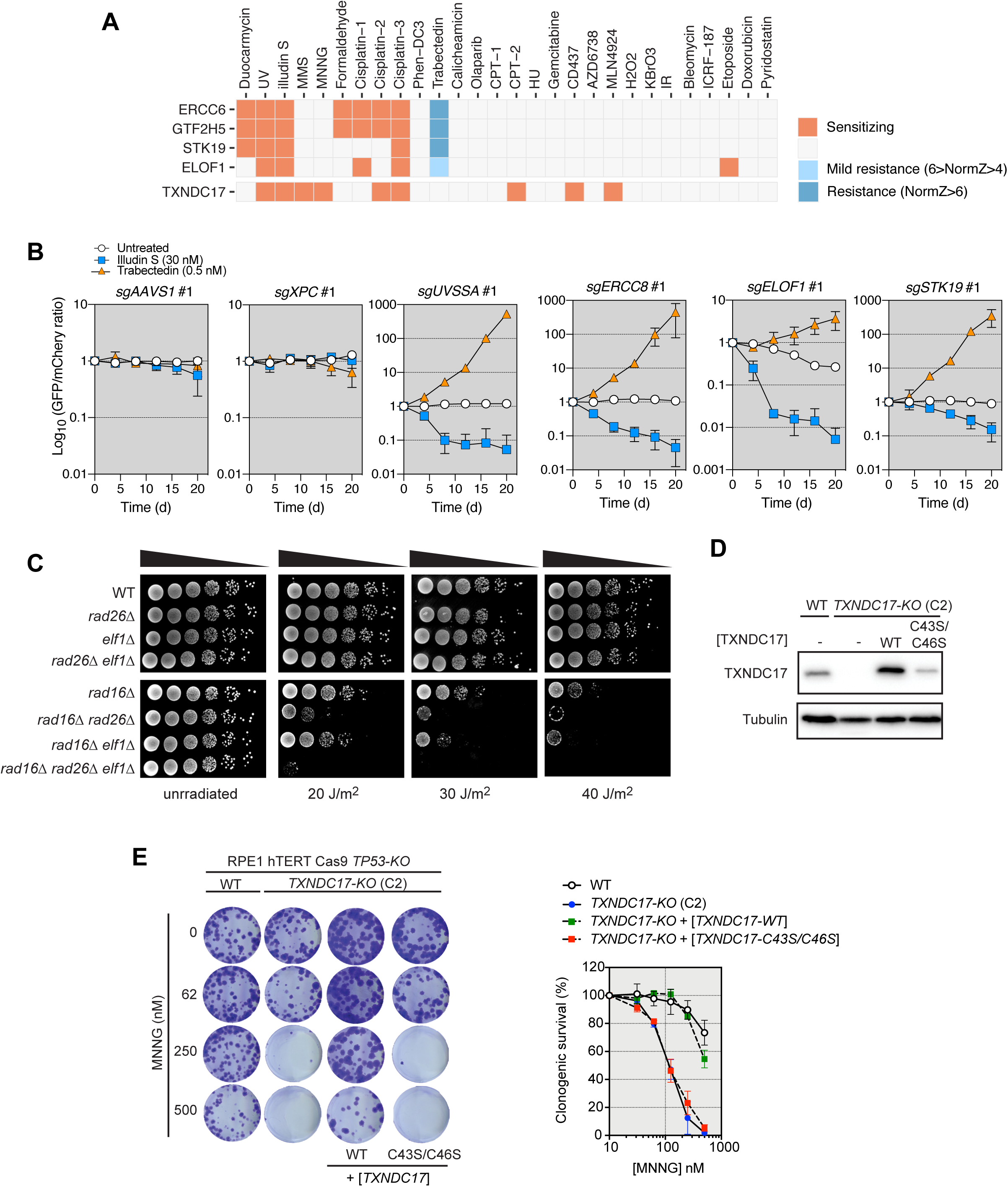
Characterization of ELOF1, STK19 and TXNDC17. (A) Fingerprint plots of *ERCC6*, *GTF2H5*, *STK19*, *ELOF1* and *TXNDC17*. The boxes are labeled according to whether mutations in these genes leads to sensitization (orange) or resistance (blue or light blue). (B) Competitive growth assays ± Illudin S (30 nM) or trabectedin (0.5 nM) treatment in RPE1 cells transduced with virus expressing the indicated sgRNAs. Data represent log_10_-transformed mean fraction of GFP-positive cells ± SD normalized to day 0 (*N*= 3, independent transductions). TIDE analysis is shown in Table S6. (C) Overnight cultures of *S. cerevisiae* strains with the indicated genotype were serially diluted and spotted onto YPD plates that were irradiated with the indicated UV dose, or left untreated. Plates were incubated at 30°C for 3 d before imaging. WT, wild type. (D) Immunoblotting for TXNDC17 in wild type (WT) or *TXNDC17-KO* cells transduced with the indicated TXNDC17-encoding viruses. Tubulin was used as a loading control. (E) Clonogenic survival of RPE1 cells of the indicated genotypes in response to increasing doses of MNNG. Representative images are shown (left) and quantified (right). Data represent mean ± SD (*N* = 3). TXNDC17 in square brackets indicates cells transfected with a wild type (WT) or mutant (C43S/C36S) expression construct. See also Figure S6.

To validate these results, we undertook competitive growth assays using illudin S and trabectedin. As positive control, we used independent sgRNAs targeting *UVSSA* and *ERCC8* (*CSA*) and as negative controls, we used guides targeting *AAVS1* and *XPC*. As expected, loss of UVSSA and CSA led to sensitization to illudin S and resistance to trabectedin (Figures 5B, S6A and Table S6). Independent sgRNAs targeting *ELOF1* similarly led to sensitization to illudin S and resistance to trabectedin in RPE1 cells, although the magnitude of the phenotypes caused by ELOF1 depletion was generally milder than those caused by depletion of UVSSA or CSA (Figures 5B, S6A and Table S7).

Given the presence of a yeast homolog of *ELOF1* in the budding yeast *Saccharomyces cerevisiae* (*ELF1*), we next assessed whether Elf1 also mediates the repair of UV-induced DNA lesions in fungi. To do so, we tested the sensitivity of an *ELF1* deletion (*elf1Δ*) alone or in combination with various mutants. In particular, we combined *elf1Δ* with a *rad16Δ* mutation that disrupts GG-NER (Prakash and Prakash, 2000). In yeast, loss of TC-NER factors only causes mild UV sensitivity on their own but their loss greatly enhances sensitivity of GG-NER mutants such as *rad16Δ.* We also prepared the *rad26Δ rad16Δ* and *rad26Δ elf1Δ* double mutants as well as the *elf1Δ rad26Δ rad16Δ* triple mutant. Rad26 is the budding yeast ortholog of CSB (ERCC6) and is involved in TC-NER. Cultures were spotted on media and were subjected to increasing doses of UV light. While the *elf1Δ* and *rad26Δ* mutants had no discernable sensitivity to UV, combining these mutations with *rad16Δ* led to an increase in UV sensitivity (Figure 5C), a phenotype that could be reversed by re-introduction of *ELF1* on a plasmid (Figure S6B). Interestingly, the *elf1Δ rad26Δ rad16Δ* triple mutant was slightly more sensitive to UV than the *rad26Δ rad16Δ* mutant indicating that *ELF1* promotes UV resistance at least partly independently of *RAD26* (Figure 5C). Given the location of Elf1 within the elongating RNA polymerase II complex, we speculate that ELOF1 may represent a previously unassigned and evolutionarily conserved modulator of transcriptional pausing in response to RNA polymerase-blocking lesions.

A second gene firmly embedded in the NER cluster was *STK19*. STK19 was first characterized as an atypical protein kinase (Gomez-Escobar et al., 1998) and *STK19* mutations are recurrent in melanoma (Hodis et al., 2012). Recent work extended these original observations and identified a role for the kinase activity of STK19 in driving NRAS-dependent melanomagenesis (Yin et al., 2019). However, STK19 was also previously reported to participate in the response to UV, in particular the transcriptional recovery following UV irradiation (Boeing et al., 2016). The presence of STK19 in the NER cluster supports the observations of Boeing and colleagues. In particular, *STK19* scored as the top illudin S-sensitizing hit and as the top trabectedin-resistance hit (Table S2). We confirmed these observations with two independent sgRNAs in competitive growth assays with both compounds (Figure 5B and Table S6). These results strongly suggest that STK19 can be assigned to the TC-NER pathway.

### TXNDC17 links redox control to genotoxin resistance

The FA/ICL cluster contains nearly all known FANC genes along with other genes coding for factors known to participate in ICL repair such as newly characterized C17orf53/HROB, which recruits the MCM8/MCM9 helicase to DNA damage sites (Hustedt et al., 2019b) (Figure 3D). Two previously uncharacterized genes in this cluster were *CCAR1* and *TXNDC17*. CCAR1 is the paralog of DBC1 (also known as CCAR2) and has been previously associated with DNA damage-induced apoptosis although these studies have not excluded a more direct role of CCAR1 in the response to DNA damage. *TXNDC17*, on the other hand has no previous link to DNA damage and *TXNDC17*-targeting sgRNAs sensitized cells to a variety of DNA damaging agents, primarily but not limited to, alkylating agents MMS and MNNG as well as cisplatin (Figure 5A and Figure S6C). TXNDC17 is a thioredoxin-domain containing protein and is one of at least 27 such proteins encoded in the human genome (Benham, 2012). Of the 25 thioredoxin-domain containing proteins screened, only a total of three showed any effect and TXNDC17 was remarkable in that it was the sole member whose loss led to strong sensitivity to more than one genotoxic agent (Figure S6C). This activity profile argues against the possibility that a non-specific redox imbalance caused by TXNDC17 depletion leads to genotoxin sensitivity, and is in line with the observation that genome maintenance mechanisms are regulated by redox sensing, such as the modulation of replication by the peroxiredoxin PRDX2 (Somyajit et al., 2017).

To validate these results, we generated independent clonal knockouts of *TXNDC17* in RPE-hTERT *p53^-/-^* cells (Figure S6D) and undertook clonogenic survival assays with MMS, MNNG, cisplatin and KBrO_3_. We observed that *TXNDC17-KO* cells are hypersensitive to MMS, MNNG and cisplatin but not to bromate, an oxidizing agent, suggesting some selectivity towards alkylating compounds (Figure S6E). The resistance to MNNG required its catalytic cysteines, suggesting that the oxidoreductase activity of TXNDC17 is involved in resistance to these genotoxins (Figure 5DE), and pointing to the possibility that the redox state of specific proteins or metabolites controls the response to DNA alkylating agents.

### CYB5R4 as a candidate modulator of protein phosphatases

*CYB5R4* (also known as *NCB5OR*), encodes another oxidoreductase that promotes the cellular resistance to multiple agents, including CPT for which it was the top hit in the CPT-2 screen (Figure 6A and Table S2). Mice lacking CYB5R4 are prone to developing diabetes (Xie et al., 2004), which made the link to *CYB5R4* genotoxin resistance intriguing. We validated that independent *CYB5R4*-targeting sgRNAs lead to CPT sensitivity in competitive growth assays (Figure 6C and Figure S6F). Interestingly, in both the DNA damage network and in a co-dependency network derived from the DepMap project (Tsherniak et al., 2017) (Figure 6B), *CYB5R4* is connected to Ser/Thr phosphatases and phosphatase regulators such as *PPP2R4* and *TIPRL*, which encode proteins that regulate the assembly and disassembly of PP2A-family holoenzymes, respectively (Guo et al., 2014; Wu et al., 2017). Like those targeting *CYB5R4*, sgRNAs against *PPP2R4* caused clear sensitization to CPT (Figures 6C, S6F and Table S6) and thus we speculate that CYB5R4 regulates the response to DNA damage indirectly via the modulation of PP2A-family phosphatases.

**Figure 6.**
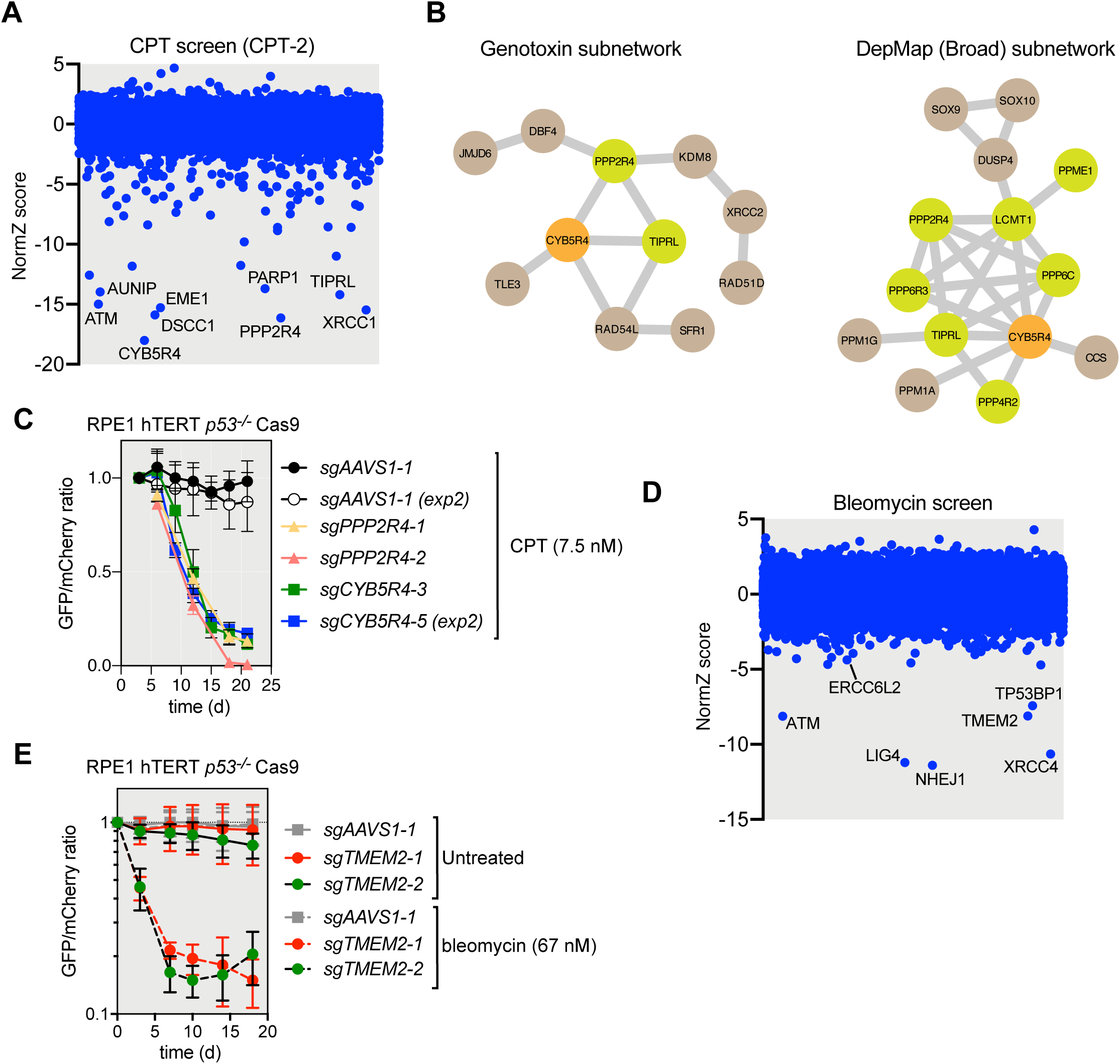
CYB5R4 and CEMIP2 promote cellular resistance to genotoxins. (A) CRISPR dropout screen results for RPE1 cells exposed to camptothecin (screen CPT-2). The 9 genes with the lowest NormZ values are labeled. (B) Subnetworks derived from the PCC genotoxin network derived in this study or from the Broad DepMap data show that CYB5R4 loss correlates with the loss of the PPP2R4 and TIPRL phosphatase biogenesis factors. (C) Competitive growth assays ± CPT (7.5 nM) in RPE1 cells transduced with virus expressing the indicated sgRNAs. Data represent mean fraction of GFP-positive cells ± SD normalized to day 0 (*N*= 3, independent transductions). TIDE analysis is shown in Table S6. (D) CRISPR dropout screen results for RPE1 cells exposed to bleomycin. The 6 genes with the lowest NormZ values are labeled. ERCC6L2 is also labeled. (E) Competitive growth assays ± bleomycin (67 nM) in RPE1 cells transduced with virus expressing the indicated sgRNAs. Data represent mean fraction of GFP-positive cells ± SEM normalized to day 0 (*N*= 3, independent transductions). TIDE analysis is shown in Table S6.

### A selection of additional genes of interest

Exploration of the individual screens and the correlation network reveals many additional notable genes. Below is a selection of those that caught our attention. One such gene is *TMEM2* (also known as *CEMIP2*) that codes for a cell surface hyaluronidase (Yamamoto et al., 2017). Its loss in the context of the screen led to a strong sensitization to the radio-mimetic chemotherapeutic bleomycin (Figure 6D), which we have confirmed in competitive growth assays with independent sgRNAs (Figure 6E). Given the cell surface localization of CEMIP2 and its biochemical activity in degrading glycosaminoglycans (Yamamoto et al., 2017), a class of polysaccharides, we speculate that TMEM2 could be an enzyme that hydrolyzes the sugar moieties of the bleomycin molecule, even though the disaccharides of glycosaminoglycans and bleomycin are chemically distinct. These results also suggest that TMEM2 levels may be a major modulator of bleomycin responses in vivo.

Formaldehyde is an environmental genotoxin that is also produced endogenously from various demethylation reactions. Formaldehyde reacts with DNA to generate DNA-DNA and DNA-protein crosslinks as well as various DNA mono-adducts (Swenberg et al., 2013), and may represent the endogenous genotoxin causing the Fanconi Anemia phenotype (Garaycoechea et al., 2018). Formaldehyde is detoxified primarily by the ADH5 enzyme in mammals (Pontel et al., 2015) and accordingly, *ADH5*-targeting sgRNAs led to formaldehyde sensitization in our screens (Figure S7A). However, the same screen also identified *ESD*, encoding an *S*-formylglutathione hydrolase, as mediating normal cellular resistance to formaldehyde treatment (Figure S7A). In a number of organisms, *S*-formylglutathione hydrolase plays a key role in a glutathione-dependent formaldehyde detoxification pathway (Chen et al., 2016) raising the possibility that a similar mechanism operates in human cells.

The *USP37* and *PHF12* genes are clustered in our network in proximity to POLE3 and POLE4, two non-essential subunits of DNA polymerase *ε* (Bellelli et al., 2018a; Bellelli et al., 2018b) (Figure S3 and Table S5). POLE3/4 promote cellular resistance to ATR inhibition (Hustedt et al., 2019a) but also to a variety of other agents that perturb DNA replication, including CPT (Figure S7B and Table S2). Loss of USP37 also causes strong sensitization to ATR inhibition and CPT in our screens. While USP37 has been previously linked to genome stability and processes influenced by DNA replication (Hernandez-Perez et al., 2016; Typas et al., 2016; Yeh et al., 2015), we speculate that USP37 may modulate the ubiquitylation state of replisome components, a finding that is supported by the observation that USP37 can be detected on nascent DNA by iPOND (Dungrawala et al., 2015) and that proximity proteomics placed USP37 in vicinity of core replisome components WDHD1 and MCM helicase subunits (Yeh et al., 2015).

Lastly, loss of the BTAF1-DRAP1 complex results in strong sensitivity to cisplatin and also to the DNA polymerase *α* inhibitor CD437 (Han et al., 2016) (Figure S7B and Table S2). BTAF1 is the human homolog of yeast Mot1 and is an inhibitor of TBP, the TATA-box binding protein (Auble and Hahn, 1993). DRAP1 is a component of NC2*α*, which stimulates the action of BTAF1 on TBP (Klejman et al., 2004). Exactly how BTAF1-DRAP1 promote resistance to cisplatin and other agents is not clear but one attractive possibility relates to the role of this complex in limiting pervasive transcription, thereby limiting the potential for transcription-replication conflicts (Xue et al., 2017).

## Discussion

In this work, we present a comprehensive and unbiased characterization of the genetic architecture of the response to DNA damaging agents in a model human cell line using CRISPR screens. Using a stringent set of criteria, these screens uncovered 840 genes (nearly 5% of the known human protein-coding genes) that influence the viability of RPE1 human epithelial cells to one or more agents that cause genotoxic stress. This number highlights the far-reaching impact of DNA damage on cellular physiology and demonstrates that the optimal response to DNA lesions often necessitates the involvement of cellular processes outside the nucleus such as autophagy or Hippo signaling, which were both enriched in our dataset (Table S3 and Figure S4). Furthermore, based on PCC values, these 840 genes form a dense network that greatly help generate biological hypotheses.

To illustrate the power of this dataset in providing new insights into the organization of the DNA damage response, we uncovered previously uncharacterized DNA repair factors, revealed unsuspected drug mechanisms-of-action, and have evidence of new drug metabolism pathways. With respect to DNA repair factors, the assignment of ERCC6L2 as a new enzyme participating in the canonical NHEJ pathway is particularly noteworthy and surprising given that end-joining reactions can be efficiently reconstituted in vitro with a set of purified components that does not include ERCC6L2 (Zhao et al., 2019). While the phenotypes caused by *ERCC6L2* mutations alone are relatively mild, our network analysis placed it firmly in the end-joining cluster, which we subsequently validated in a number of assays. It remains unclear what role an enzyme of the SWI/SNF family may have during the end-joining reaction. Given that this enzyme family functions as dsDNA translocases, one simple model may be that it removes roadblocks that impair the completion of the end-joining reaction. These roadblocks could consist of nucleosomes, other DNA-bound proteins or even unproductive NHEJ assemblies that preclude completion of ligation. Understanding the role of ERCC6L2 in NHEJ will require that we understand both the biochemical activities of ERCC6L2 and the type of end-joining reaction substrates that necessitate the action of ERCC6L2. Our results also suggest that DSBs of unknown origin underlie the bone marrow failure in individuals with *ERCC6L2* mutations. Therefore, finding the endogenous sources of DSBs in the hematopoietic stem cell compartment, similarly to what was done recently for Fanconi Anemia (Pontel et al., 2015) may provide important clues as to the pathogenesis of this syndrome and may reveal paths of intervention that could benefit those patients.

In addition to identifying new DNA repair factors and new biology in the response to genotoxic stress, this work also helps solidify a number of previous observations. A case in point is the homologous recombination gene cluster, which lies at the core of our human DNA damage network. A number of genes are embedded in this cluster for which evidence of a role in homologous recombination was either circumstantial, indirect or supported by very few studies. Examples are *AUNIP, KDM8/JMJD5* and *JMJD6* (Figure S3) (Amendola et al., 2017; Huo et al., 2019; Lou et al., 2017). In particular, recent work on KDM8 and JMJD6 suggest that instead of acting as lysine demethylases they are rather acting as lysyl- and arginyl-hydroxylases, respectively (Webby et al., 2009; Wilkins et al., 2018). Whether or not this type of post-translational modification regulates HR remains to be determined but it represents a fascinating possibility.

Despite identifying nearly 5% of the human protein-coding genes as being involved in the response to DNA damaging agents, this list only represents a partial view of the architecture of the human DNA damage response. Indeed, there are inherent technical limitations of dropout chemogenomic CRISPR screens that preclude having a complete view of the genes surveyed. One such limitation is that dropout screens comparing two conditions (control vs perturbation) lack power to probe the function of many cell-essential genes. As an example of this phenomenon, the sgRNAs targeting the gene coding for the recombinase RAD51 are depleted too rapidly in our cell lines (with and without drugs) to produce statistically reliable results in most of our screens. Furthermore, our screens only used a single cell line model, RPE1-hTERT, and previous work on ATR and PARP inhibitors, among others, have revealed substantial cell line-specific responses to these drugs that are likely reflecting their biology and tissues of origin (Hustedt et al., 2019a; Zimmermann et al., 2018). Finally, we have not explored the full range of genotoxic agents and this dataset will benefit from additional screens probing the response to compounds such as antimetabolites or environmental carcinogens.

One class of genome maintenance regulators that was largely and conspicuously absent from our dataset were the modulators of DNA replication fork reversal and fork protection such as SMARCAL1, ZRANB3, HLTF and RADX (Quinet et al., 2017). An exception to this list was BOD1L, which was a prominent hit in multiple screens and clustered close to ATR and the 9-1-1 complex in our network, in line with its proposed function at replication forks (Higgs et al., 2015) (Figure S3). This relative lack of replication fork “quality control” factors is highly surprising given the current interest in fork reversal and nascent DNA degradation. These observations suggest either a high degree of genetic redundancy in this process or that these genes do not participate in the cellular resistance to the type of replication challenges that were induced by the agents screened in this study.

The work presented here has clear implications for the development of therapeutics and genetic toxicology. For example, the characterization of compounds using chemogenomic CRISPR screens or via the use of isogenic cell line panels derived from information produced in this work may enable the rapid identification of desired (or undesired) genotoxic effects such as formation of DNA double-strand breaks, DNA adducts or interference with transcription. Furthermore, the dataset presented can be mined for the development of patient-selection hypotheses as a means to identify subsets of tumors sensitive to a certain agent. For example, deep deletions and mutations in the *MCPH1* gene occurs frequently in a number of cancer types such as bladder and colorectal cancer (Figure S7C). Loss of MCPH1, a protein with dual function at centrosomes and as a poorly characterized reader of histone H2AX phosphorylation (Liu et al., 2016), leads to sensitization to multiple genotoxic drugs but scored particularly highly in the screen probing the response to etoposide, a cancer chemotherapeutic (Table S2). These results suggest that tumors with *MCPH1* loss may be particularly vulnerable to etoposide treatment, a drug already used in the clinic. Finally, our data indicates that a subset of G quadruplex ligands have the unsuspected property of trapping TOP2 in a manner that produces signaling-competent DSBs at a slow rate. Since this mechanism of action is clearly distinct from canonical TOP2 poisons like etoposide, it is conceivable that the TOP2-trapping property of these agents could be exploited therapeutically.

## Methods

### Cell lines

Generation of RPE1-hTERT Cas9 *TP53-KO* (RPE1) has been previously described (Zimmermann et al., 2018). Cas9-expressing cell lines were maintained in the presence of 2 µg/mL blasticidin. RPE1 *TDP2^-/-^* cells were generated by transfecting with the sgRNA targeting *TDP2* (see Table S6) using RNAiMAX (Invitrogen) according to manufacturer’s protocol. 24 h after transfection, cells were split and seeded in a 60 mm plate. After an additional 72 h, single cells were sorted into 96-well plates on a BD FACSAria Cell Sorter instrument and grown until colonies formed. *TDP2^-/-^* clones were selected on the basis of successful gene editing determined by PCR amplification and TIDE analysis (Brinkman et al., 2014).

sgRNAs used for transfection were ordered from Integrated DNA Technologies. crRNAs containing the target-specific sequence for guiding Cas9 protein to a genomic location were annealed with tracrRNA to form a functional sgRNA duplex. DT40 knockout constructs for *ERCC6L2*, *XRCC4* and *XLF* were generated as previously described using a MultiSite Gateway three-fragment vector construction kit. 5’ and 3’ arms were cloned into the pDONR P4-P1R and pDONR P2R-P3 vector, respectively. The knockout constructs were generated by attL×attR recombination of the pDONR-5′ arm, pDONR-3′ arm, resistant gene cassette-containing pDONR-211 and the pDEST R4-R3 destination vector. *TXNDC17* gene knockouts were generated by electroporation of LentiGuide vectors (see Table S6) using an Amaxa II Nucleofector (Lonza, Basel). 24 h after transfection, cells were selected for 48 h with 15 µg/mL puromycin, followed by single clone isolation. Gene mutations were further confirmed by PCR amplification and TIDE analysis. Stable integration of pHIV-TXNDC17 in RPE1-hTERT *TP53^-/-^* cells was achieved by lentiviral transduction. Transduced RPE1 cells were selected by culturing in the presence of 400 µg/ml nourseothricin (Jena Bioscience, cat # AB-102L). Anti-TRP14 antibody (1:1,000, Abcam #ab121725) was used to validate correct knock-out and complementation of TXNDC17. CH12F3-2 murine B cell lymphoma mutant lines were edited by transient transfection with pX330 plasmid constructs expressing sgRNAs against *Trp53bp1* and *Ercc6l2* and were selected on the basis of successful gene editing determined by PCR amplification and TIDE analysis (Brinkman et al., 2014) (see Table S6). HEK293F cells (Invitrogen) were cultured in DMEM with 10% (v/v) fetal bovine serum (Gibco), 50 U/µL penicillin, 50 µg/mL streptomycin, and 0.5 mmol/L sodium pyruvate at 37°C in 5% CO_2_atmosphere.

### Cell culture

RPE1-hTERT and 293T cells were purchased from ATCC. U2OS EJ7 cells originate from the laboratory of Jeremy Stark and were a kind gift. RPE1-hTERT-derived cell lines, 293T and derived cell lines were grown in Dulbecco’s Modified Eagle Medium (DMEM; Gibco/Thermo Fisher #11965092) supplemented with 10% fetal bovine serum (FBS; Wisent), 1x GlutaMAX, 1× non-essential amino acids (both Gibco/Thermo Fisher), 100 U/mL penicillin and 100 μg/mL streptomycin (Pen/Strep; Wisent). U2OS-derived cell lines were grown in McCoy’s 5A (Gibco #1660-082) + 10% FBS (Gibco) + 100 U/mL penicillin and 100 μg/mL streptomycin (Pen/Strep; Wisent) + 1x L-Glutamine (final 2 mM, Wako Chemical #073-05391). DT40 cells were grown in RPMI-1640 medium + 10% FBS (Gibco) + 2% Chicken serum (Gibco) + 100 U/mL penicillin and 100 μg/mL streptomycin (Pen/Strep; Wisent) + 10 mM HEPES and grown at 39°C. All other cell lines were grown at 37°C and atmospheric O_2_. All cell lines were routinely authenticated by STR and tested negative for mycoplasma. The two-color competitive growth assays were performed as previously described (Hustedt et al., 2019a). Imaging was done on an IN Cell Analyzer system (GE Healthcare Life Sciences) equipped with a 4X objective. Segmentation and counting of GFP- and mCherry-positive cells was performed using an Acapella script (PerkinElmer).

### CRISPR screens

RPE1-hTERT Cas9 *TP53-KO* cells were transduced with the lentiviral TKOv2 or TKOv3 library (Hart et al., 2015; Hart et al., 2017) at a low MOI (∼0.35) and puromycin-containing medium was added the next day to select for transductants (see Table S1 for more details). The next day cells were trypsinized and replated in the same plates while maintaining the puromycin selection. 3 days after infection, which was considered the initial time point (t0), cells were pooled together and divided in 2 technical replicates (usually labelled as “A” and “B”). Negative-selection screens were performed by subculturing cells at days 3 and 6 (t3 and t6), at which point each replicate was divided into different treatments, including one that was left untreated as a control (NT), and then subcultured every 3 days (t9, t12 and t15) until the final timepoint at t18. For all screens, drugs and treatments were applied according to previous evaluation of LD20 concentrations in uninfected RPE1 cells treated for 12 days (except for gemcitabine, which was determined over 3d). Cells were subcultured and medium with and without drugs was refreshed every 3 days. UV and IR treatments were applied one day after subculturing cells (t7, t10, t13 and t16). Cell pellets were frozen at day 18 for gDNA isolation. Screens were performed in technical duplicates and library coverage of ≥ 375 cells per sgRNA was maintained at every step with the exception of the gemcitabine screen which was carried out a 250-fold coverage. gDNA from cell pellets was isolated using the QIAamp Blood Maxi Kit (Qiagen) and genome-integrated sgRNA sequences were amplified by PCR using Q5 Mastermix (New England Biolabs Next UltraII, M5044S). i5 and i7 multiplexing barcodes were added in a second round of PCR and final gel-purified products were sequenced on Illumina NextSeq500 systems to determine sgRNA representation in each sample.

### Functional Enrichment Analysis

g:Profiler (https://biit.cs.ut.ee/gprofiler/gost) was used to identify functional enrichment analysis of the 840 hit genes (NormZ < −3 and FDR < 15% for sensitizing genes; NormZ > 6 for resistance genes), resulting in a list of 1230 statistically significant enriched terms (Benjamini-Hochberg FDR < 0.05). The hit genes list was treated as an unordered query, and statistical tests were conducted within a statistical domain scope of only annotated genes, selecting terms with sizes between 2 and 500 genes, considering the GO molecular function, GO cellular component, GO biological process, KEGG, Reactome and WikiPathways data sources, removing electronic GO annotations. The Ensembl ID with the most GO annotations were chosen for all 5 ambiguous genes (*BABAM1*, *CHTF8*, *FOXP1*, *MATR3*, *SOD2*).

### DNA repair-associated terms gene list

The list of 336 genes considered as ‘DNA repair genes’ was assembled by combining all genes present in at least one of 100 manually selected terms present in the g:Profiler data sources mentioned under ‘Functional Enrichment Analysis’. The terms were selected by their association with DNA damage and repair such as Homologous recombination (KEGG:03440), DNA Double-Strand Break Repair (REAC:R-HSA-5693532), response to UV (GO:0009411); DNA damage response (WP:WP707) and negative regulation of cell cycle phase transition (GO:1901988).

### Network Analysis

The Network Analysis can be divided in two phases: the exploratory phase and the design phase. For the exploratory phase we developed an R shiny app (https://hs-durocherlab.shinyapps.io/ScreenClustering/) that would allow us to visualize several distance measures, clustering techniques and networks graphs, with dynamically changing parameters to provide insights into our research. The design phase consisted of choosing how best to illustrate a single meaningful network of the 840 gene hits. For this step the interaction network was created using Cytoscape version 3.7.1, where the 568 nodes represent genes and the 1231 edges represent all pairwise connections that had a Pearson’s correlation coefficient (PCC) greater than 0.725, explaining why the network has fewer nodes than the initial 840 genes. These arbitrary threshold values were chosen as such to ensure that we captured interesting known genetic interactions but could filter out uncertain values and be able to observe some structure in the network. The PCC for each pair of genes was calculated using the default parameters of function ‘*cor’*, from the *stats* package, R version 3.5.2, from our data frame of 840 genes, each with 25 NormZ scores. Data from Cisplatin-1, Cisplatin-2 and CPT-1 screens were omitted when calculating the PCC to avoid skewness towards a specific drug.

### Radar and Fingerprint Plots

Radar charts were drawn in R version 3.5.2, using the *fmsb* package version 0.6.3, treating each screen independently. The one-sided alternative hypothesis Fisher Exact Test p-value was calculated by testing the enrichment [FDR < 15% for sensitizing genes and FDR < 1% for resistance genes] of each single pathway (Table S4) individually against the expected enrichment for that screen excluding that pathway, using the *‘fisher.test’* function from the *stats* package, version 3.5.2. The Bonferroni correction threshold, adjusted for the number of pathways tested [for 9 pathways, and *α* = 0.05, the threshold was therefore -log2(0.05/9) *≅* 7.49], was also added to the graph. Fingerprint plots were drawn using the *ggplot2* package version 3.2.0.

### Data Quality Control

For each screen, at least two replicates of each condition were performed and to assess data quality, the two main data diagnostic tools used were the MAGeCK (Li et al., 2014)standard count summary R script and the BAGEL precision-recall curves using the core essential (CEGv2.txt) and nonessential (NEGv1.txt) gene lists from (https://github.com/hart-lab/bagel). From the MAGeCK count summary we observed that most samples had a mapped reads percentage above 78%, with only calicheamicin replicate B (67%) and trabectedin replicate B (68%) having mapped reads percentages below 70%. Gini indexes were acceptably low, with most screens scoring below 0.15; only camptothecin-1 replicate B had a high Gini index of 0.21 in comparison. From the precision-recall curves it was possible to verify that known essential and nonessential genes behaved as expected, even though some samples had a steeper curve than most others.

### EJ7 assay

U2OS-EJ7 cells were infected with viruses produced in HEK293T using second generation vectors psPAX2 (Addgene #12259) as packaging plasmid, pMD2.G (Addgene #12259) as envelope and pLentiCRISPRv2-mCherry (Addgene #99154) as transfer vectors. All sgRNAs cloned into pLentiCRISPRv2-mCherry are reported in Table S6. After 6 d in culture, cells were plated at 70-80% confluence in 6 well plates and transfected with 1μg of each plasmid containing 7a and 7b sgRNAs (Addgene #113620 and #113624) using Lipofectamine 2000 (ThermoFisher, #11668030). 48 h after transfection cells were resuspended, washed in PBS and analysed using a BD LSR Fortessa.

### High content imaging

Cells were seeded (∼10,000 cells/well) in 96-well plates and cultured for 24 h. 20 μmol EdU (5-ethynyl-2-deoxyuridine, Life Technologies) was added 30 min prior washing with PBS and fixation with 4% paraformaldehyde (PFA) in PBS for 10 min. Cells were rinsed with PBS and permeabilized using 0.3% TritonX-100/ PBS for 30 min. Cells were washed with PBS and incubated in blocking buffer (0.2% fish skin gelatin, 0.5% BSA/PBS) for 30 min. Fresh blocking buffer containing mouse anti-γH2AX (Millipore #JBW301, 1:5,000) was added for 2 h. Cells were rinsed three times with PBS and blocking buffer with AlexaFluor 488-coupled goat anti-mouse antibody (Life Technologies, 1:1000) and 0.8 μg/mL DAPI (4,6-diamidino-2-phenylindole, Sigma) was added for 1h. After rinsing with PBS, immunocomplexes were fixed again using 4% PFA/PBS for 5 min. Cells were rinsed with PBS and incubated with EdU staining buffer (150 mM Tris/HCl pH8.8, 1mM CuSO4, 100 mM ascorbic acid and 10 μM AlexaFluor azide (Life Technologies)) for 30 min. After rinsing with PBS, images were acquired on an In Cell Analyzer 6000 automated microscope (GE Life Sciences) with a 60X objective. Image analysis was performed using Columbus (PerkinElmer). Cell cycle profiling and analysis was evaluated based on EdU and DAPI staining.

### RADAR assay for detection of TOP2 cleavage complexes

The assay was performed as previously described in (Anand et al., 2018). RPE1-hTERT Cas9 *TP53-KO* cells were seeded in 6 well plates (1×10^6^ cells each). 24 h later cells were lysed directly in the plates using 1 mL of buffer containing 5 M guanidinium isothiocyanate, 10 mM Tris (pH 6.5), 20 mM EDTA, 4% TritonX-100/PBS, 1% Sarkosyl (Sigma-Aldrich #L9150), 1% DTT. Genomic DNA was precipitated incubating the lysate at −20°C for 5 min with 0.5 volumes of 100% ethanol, shaking vigorously, and centrifuging at 14,000 rpm for 15 min at 4 °C. The DNA pellet was washed 2 times in 75% EtOH by vortexing and centrifuging again at 14,000 rpm for 15 min at 4 °C. The final DNA pellet was resuspended in 100 µL of in freshly prepared 8 mM NaOH solution. The samples were incubated at 65°C for 5 min, passed through a 26G needle 10 times and then quantified using a Nanodrop (ThermoFisher). For each sample, 7 µg of DNA was resuspended in 200 µL of 25 mM NaPO_4_ (pH 6.5) buffer and loaded in a slot blot apparatus (Bio-Dot® SF Microfiltration Apparatus, Bio-Rad). Membranes were blocked for 1 h in 5% milk/TBS Tween-20 solution and incubated for 1 h with anti-TOP2β (1:5,000, mouse anti-human, Clone 40/Topo IIβ, RUO – 611492, BD Biosciences-US) or anti-TOP2α (1:10,000, mouse anti-human, Topo IIα antibody (F-12), sc-365916, Santa Cruz) diluted in 5% milk/TBS Tween-20.

### Class switch recombination assays

To induce switching in CH12F3-2 murine B cell lymphoma cell or their derivatives, 2×10^5^ cells were cultured in CH12 medium supplemented with a mixture of IL4 (10 ng/mL, R&D Systems #404-ML-050, Minneapolis, MN, USA), TGFβ (1 ng/mL, R&D Systems #7666-MB-005) and anti-CD40 antibody (1 μg/mL, #16-0401-86, eBioscience, Thermo Fisher) for 48 h. Cells were then stained with anti-IgA-PE and the fluorescence signal was acquired on an LSR II or Fortessa X-20 flow cytometer (BD Biosciences). AID protein was detected by immunoblotting with mouse mAb L7E7 (Cell Signaling #4975) and normalised to β-actin (Sigma #A2066) levels. Quantitative PCR was performed with qPCRBIO SyGreen Blue mix (PCR Biosystems) and CFX384 Real-Time PCR Detection System (Bio-Rad) according to manufacturers’ instructions. The Iμ sterile transcript was detected with primers 5’-gaacatgctggttggtggtt-3’ and 5’-tcacacagagcatgtggact-3’; the Iα transcript was detected with primers 5’-gggacaagagtctgcgagaa-3’ and 5’-tcaggcagccgattatcact-3’, and normalised to HPRT (primers 5’-cccagcgtcgtgattagc-3’ and 5’-ggaataaacactttttccaaat-3’). All samples were collected 48 h after stimulation with anti-CD40, IL4, and TGFβ as described above.

### Clonogenic survival assays

RPE1 and RPE *TXNDC17-KO* cells were seeded in 6-well plates (250 cells per well) and treated with indicated concentrations of MMS, cisplatin, MNNG, or KBrO3. After 14 days, colonies were stained with crystal violet solution (0.4% (w/v) crystal violet, 20% methanol) and manually counted. Relative survival was calculated for the drug treatments by setting the number of colonies in non-treated controls at 100%.

### Plasmids and viral vectors

DNA corresponding to sgRNAs was cloned into LentiGuide-Puro (Addgene: 52963), LentiCRISPRv2 (Addgene: 52961), or a modified form of LentiGuide-Puro in which Cas9 was replaced by NLS-tagged GFP or mCherry using AgeI and BamHI (designated as LentiGuide-NLS– GFP or –mCherry respectively), as described (Noordermeer et al., 2018). Lentiviral particles were produced in 293T cells by co-transfection of the targeting vector with vectors expressing VSV-G, RRE and REV using calcium-phosphate, PEI (Sigma-Aldrich) or Lipofectamine 2000 (ThermoFisher). Viral transductions were performed in the presence of 8 µg/mL polybrene (Sigma-Aldrich) at an MOI <1, unless stated otherwise. Transduced RPE1 cells were selected by culturing in the presence of 20 µg/mL puromycin for 3 d. The TXNDC17 coding sequence was obtained from the ORFeome collection (http://horfdb.dfci.harvard.edu/), archived in the Lunenfeld-Tanenbaum Research Institute’s OpenFreezer (Olhovsky et al., 2011). The complete TXNDC17 coding sequence was PCR-amplified and cloned into the pHIV-NAT-T2A-hCD52 vector using NotI and EcoRI restriction enzymes. To generate a catalytic dead TXNDC17 mutant, the active-site Cys residues were replaced by Ser (C43/46S) using PCR-mediated site-directed mutagenesis.

### Cell proliferation assays

RPE1 cells were seeded in 96-well plates (2,000 cells per well) and treated with sequential serial dilutions of genotoxic agents as indicated. After 4 days of treatment, the cell confluency was measured using an IncuCyte Live-Cell Analysis (Sartorius). Confluence growth inhibition was calculated as the relative confluency compared to untreated cells.

### YFP-TOP2α overexpression and pyridostatin sensitivity

To generate cells that overexpress YFP–TOP2α, HEK293F were transfected with a plasmid encoding YFP-TOP2α (Schellenberg et al., 2018), followed by growth under selection with 20 μg/mL blasticidin for 3 weeks. Cells with high levels of YFP expression were selected by FACS. A single YFP-positive cell was propagated to generate the cell line that overexpresses YFP–TOP2α. 96-well plates were seeded with 2000 cells per well of the indicated cell line in 200 μL DMEM media. After growth for 24 h, the indicated concentration of DNA damaging agent was added. For proliferation (AUC) assays, cells were grown in an IncuCyte Zoom live cell imager (ESSEN Biosciences) with images recorded every 2 h for 4 d after addition of drug. The confluence percentage for each image was calculated using the IncuCyte Zoom software and summed over the course of the experiment for each well. Viability measurements were normalized to that of untreated cells and fit to a 4-parameter dose-response curve to calculate IC_50_ values.

### Yeast strains and spot assay

The *S. cerevisiae* strains in used this study are all derivatives of S288c BY4741 (Brachmann et al., 1998) and are described in Table S7. BY4741-derived *elf1::KANMX* and *rad26::KANMX* deletion strains as well as a Y7092-derived *rad16::NATMX* strain were kind gifts of Charlie Boone and were used as starting strains or sources of disrupted alleles. We constructed all strains using standard genetic techniques. The original *elf1::KANMX* strain we received was discovered to have unusual growth properties on synthetic media, therefore we recreated the *elf1* deletion strain by amplifying the *elf1::KANMX* genomic locus by PCR, transforming the reaction product into *ELF1* wild-type yeast and screening G418-resistant colonies for normal growth rates and by diagnostic PCR for correct integration.

*ELF1* rescue plasmids were created by ligating PCR products from wild-type genomic DNA template into the BamHI/XhoI sites of either pRS416 to insert the *ELF1* locus (including the open reading frame with 133 and 109 bp of up- and downstream flanking genomic DNA, respectively), or into a derivative of pRS416-ADH where the *ELF1* open reading frame was placed under control of the *ADH1* promoter.

To measure UV sensitivity, cells were grown at 30°C overnight in 2% glucose-containing non-selective rich (XY) media or synthetic complete media lacking uracil to maintain plasmid selection. The next day serial ten-fold dilutions of cells were spotted on XY plates and exposed to the indicated dose of UV treatment. Plates were imaged after 3 days of incubation at 30°C in the dark.

## Acknowledgments

We thank Jing Yi Yuan for technical assistance on some of the screens. We are indebted to Jason Moffat for the TKOv2 and v3 libraries and to Stephane Angers and Traver Hart for developing tools and protocols for CRISPR screening in Toronto and to Charlie Boone for yeast strains. We also thank Feilong Meng, Sven Rottenberg and Martijn Luijsterburg for sharing unpublished results. AAQ, GSM and AMH are recipients of long-term EMBO fellowships (ALTF 910-2017, 795-2017 and 124-2019, respectively), NH was supported by a Human Frontier Science Program long-term Fellowship, SER is supported by a fellowship from AIRC and SA was a Banting post-doctoral fellow. ASB was supported by a PhD Studentship from AEFAT (Asociación Española Familia Ataxia Telangiectasia) and an EMBO short-term fellowship for a visit to the DD laboratory. The ICRF187 screen in FCL laboratory was funded by grants from the Spanish Government (SAF2017-89619-R, European Regional Development Fund) and the European Research Council (ERC-CoG-2014-647359). Work in the RSW laboratory was supported in part by the US National Institute of Health Intramural Program, US National Institute of Environmental Health Sciences (NIEHS, 1Z01ES102765). DD is a Canada Research Chair (Tier I) and the work was supported from grants from the CIHR (FDN143343 to DD; FRN 123518, PJT-156330 to AM) Canadian Cancer Society grant 705644 (to DD) with additional support to DD from the Krembil Foundation.

## Figure legends

**Figure S1.**
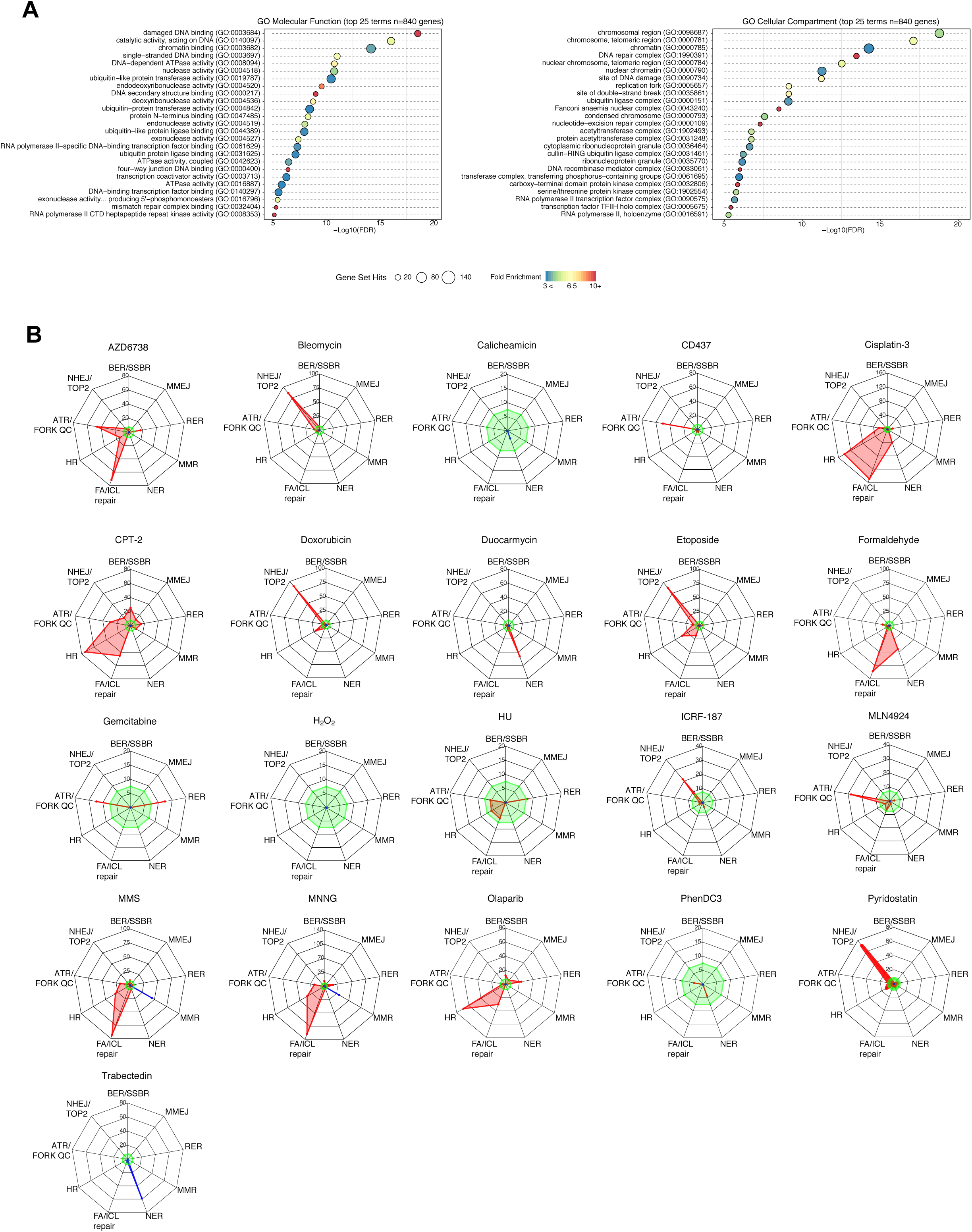
Gene term and DNA repair pathway enrichments. Relates to Figure 1. (A) Top 25 enriched GO terms, for molecular function (left) and cellular compartment (right), identified using g:Profiler (>10-fold enrichment; P < 0.05, with Benjamini-Hochberg FDR correction) among the 840 gene hits. Circle size indicates the number of genes from the common gene hit list included in each GO term, x-axis position indicates -log_10_-transformed FDR score and the colors indicate the fold enrichment compared with the whole-genome reference set. (B) Radar plots of the genotoxic CRISPR screen performed with genotoxic agents not shown in Figure 1, depicting sensitization (red) or resistance (blue) for different DNA repair pathways (axes). Values indicate the log_2_-transformed p-values of the Fisher’s exact test score. Bonferroni thresholds are indicated in green.

**Figure S2.**
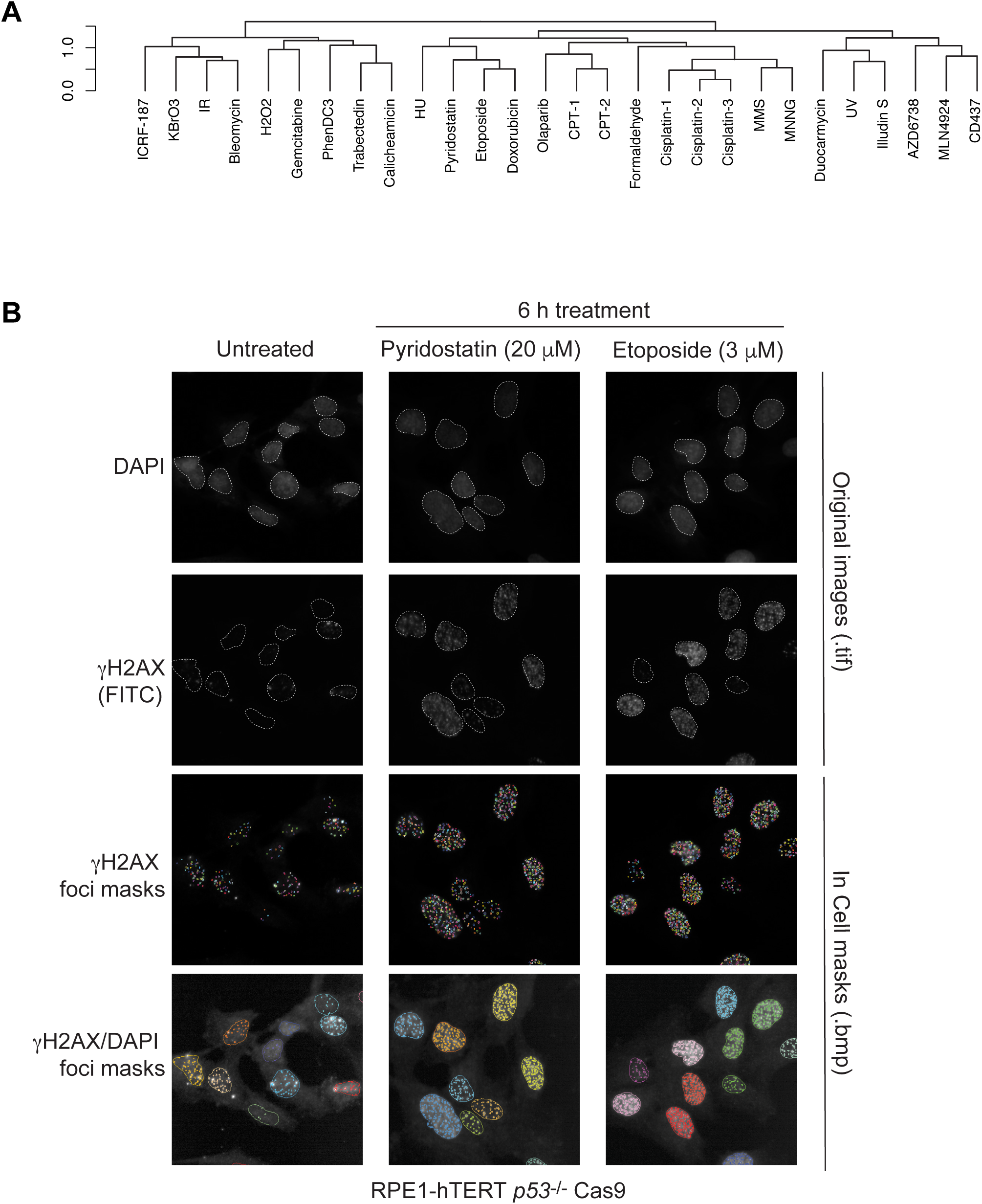
Clustering of screens and control images. Relates to Figure 2. (A) Hierarchical cluster of the 28 genotoxic CRISPR screens based on PCC values calculated using the 840 genes hits. (B) Representative images depicting automated quantation of γH2AX foci using the IN Cell Analyzer 6000 coupled with Columbus image data analysis system. The initial nuclear masks were generated using images acquired from the DAPI channel. Upon selection of only completely in-frame nuclei, the number of foci and γH2AX intensity was evaluated in the FITC (*γ*-H2AX) channel (additional details in Methods).

**Figure S3.**
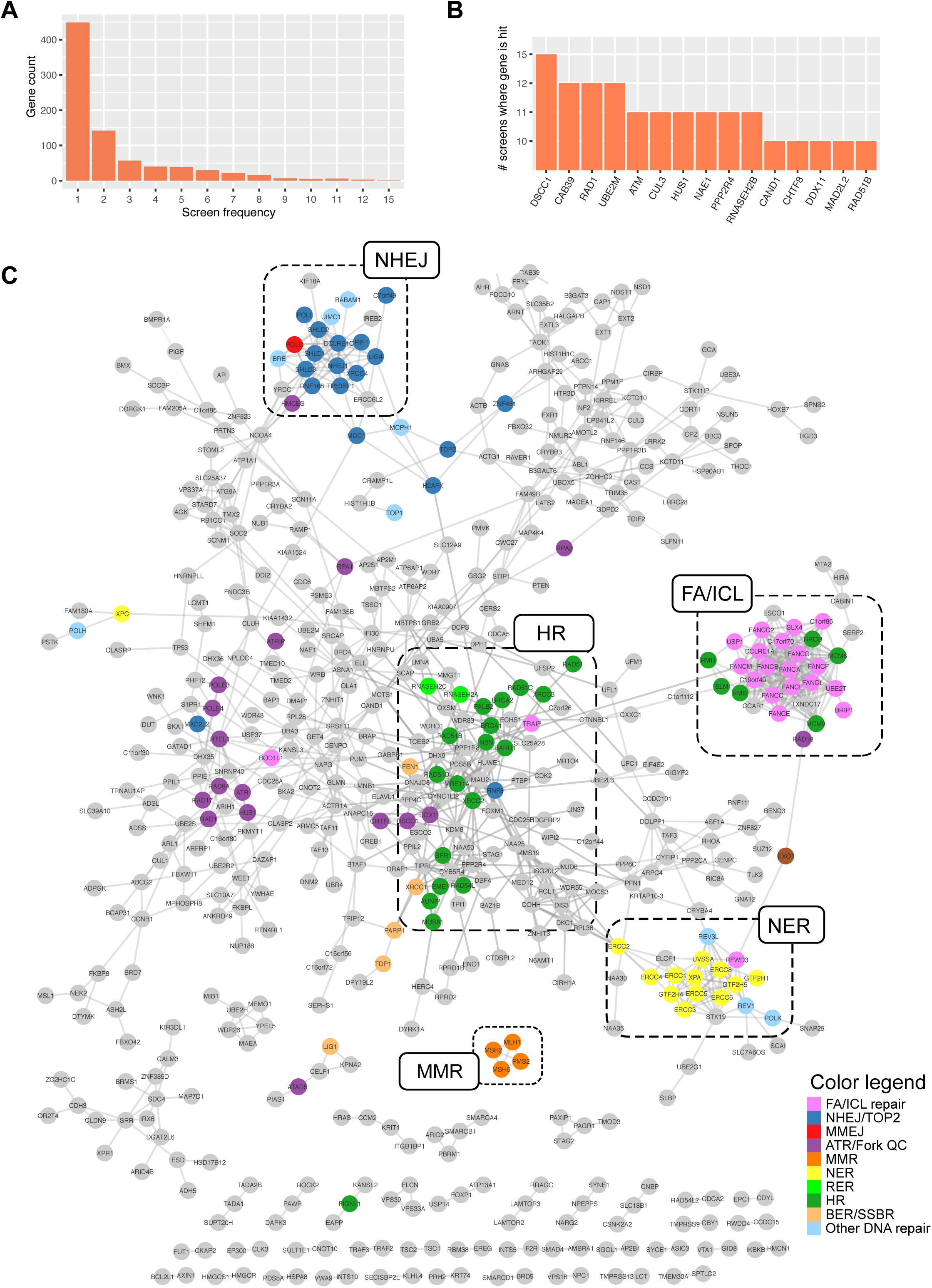
The DNA damage network with gene names. Relates to Figure 3. (A) Histogram that bins the number of genes and how many times they were identified as hits in the screens. (B) Histogram of the 10 genes that scored as hits in the highest number of screens. (C) The same PCC-based genetic network as in Figure 3 with gene names annontated.

**Figure S4.**
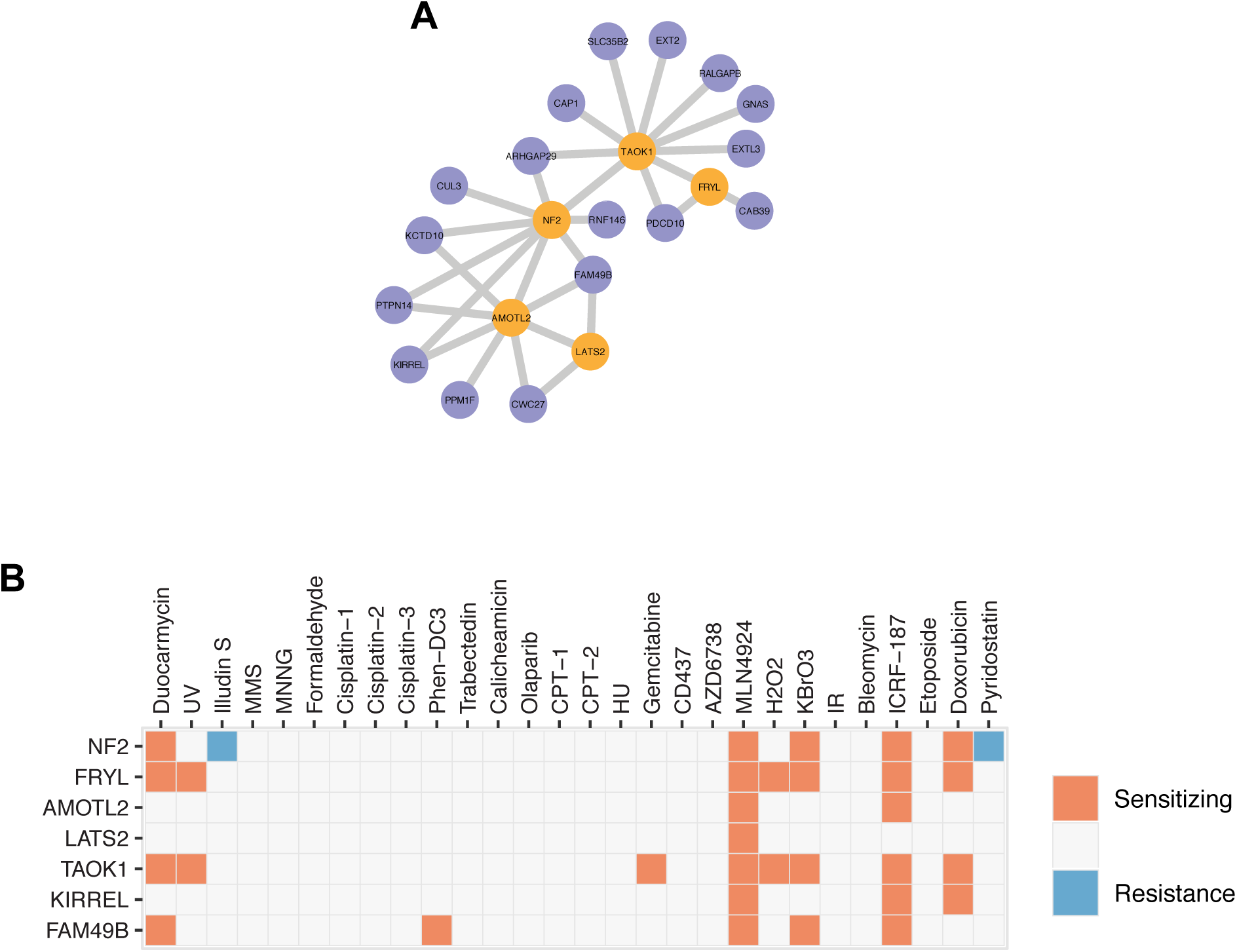
Hippo signaling promotes resistance to DNA damaging agents. Relates to Figure 3. (A) Subnetwork highlighting *NF2*, *FRYL*, *AMOTL2*, *LATS2*, *TAOK1* with genes that have PCC values > 0.725. The network was generated with Cytoscape (v3.7.1) using perfused force directed layout. (B) Fingerprint plots of *NF2*, *FRYL*, *AMOTL2*, *LATS2*, *KIRREL*, *FAM49B* and *TAOK1*. The boxes are colored according to whether mutations in these genes leads to sensitization (orange) or resistance (blue) in any of the 28 CRISPR screens.

**Figure S5.**
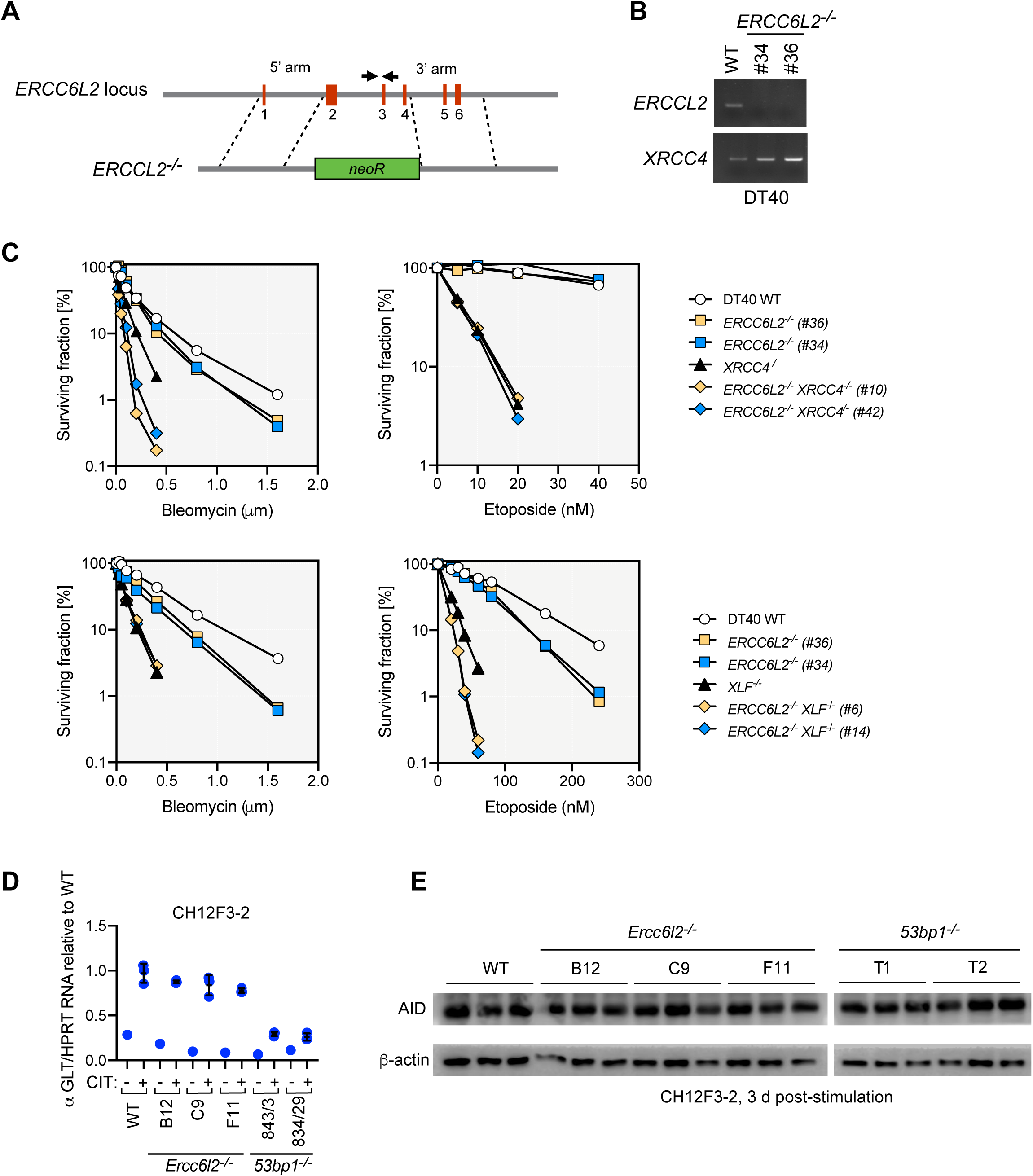
Additional characterization of ERCC6L2. Relates to Figure 4. (A) Schematic of the strategy used to knockout *ERCC6L2* in DT40 cells. Shown are the location of primers (arrows) used to confirm targeting. (B) Agarose gel images of the PCR-mediated amplification of the internal *ERCC6L2* sequence to confirm gene-targeting of *ERCC6L2*, using the primers indicated in (A). (C) Cell proliferation assays of DT40 cells of the indicated genotypes treated with either etoposide or bleomycin for a 3 d period. Data is presented as the mean of a technical triplicate. An independent experiment is shown in Figure 4. (D) Quantitative PCR assessing the levels of Iμ sterile transcript and Iα transcript (normalised to *HPRT*) in CH12F3-2 cell lines of the indicated genotypes. CIT is the cytokine cocktail used to activate switching (anti-CD40, IL-4, and TGFβ). (E) Immunoblotting of AID in different CH12F3-2 cells of the indicated genotypes. Samples were collected 3 d after stimulation (N = 3). β-actin was used as loading control.

**Figure S6.**
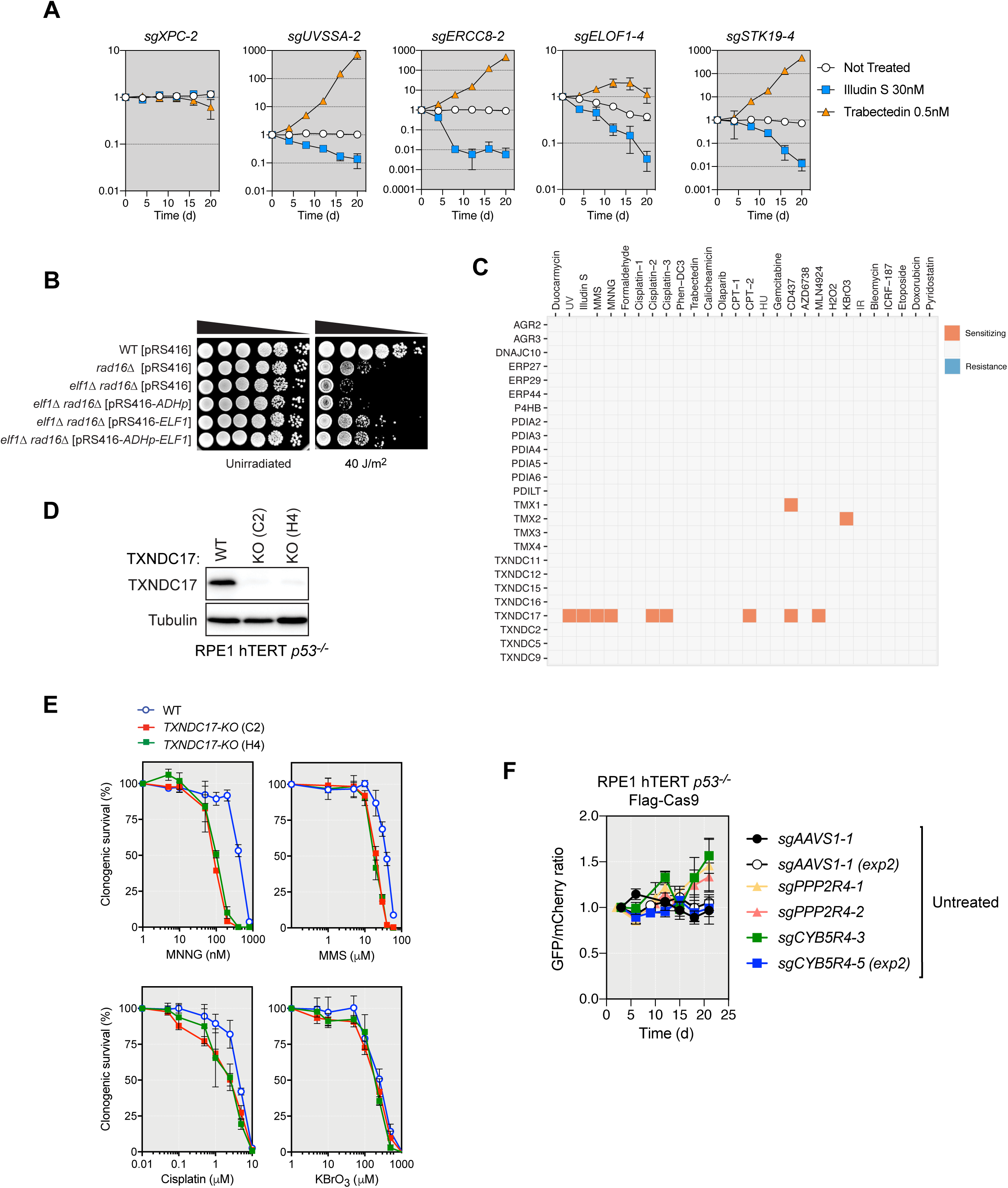
Further characterization of ELOF1, STK19, TXNDC17 and CYB5R4. Relates to Figures 5 and 6. (A) Competitive growth assays ± Illudin S (30 nM) or trabectedin (0.5 nM) in RPE1 cells transduced with virus expressing the indicated sgRNAs. Data represent log_10_-transformed mean fraction of GFP-positive cells ± SD normalized to day 0 (*N*= 3, independent transductions). TIDE analysis is shown in Table S6. (B) Overnight cultures of *S. cerevisiae* strains with the indicated genotype and transformed with the indicated plasmids were serially diluted and spotted onto YPD plates that were irradiated with the indicated UV dose, or left untreated. Plates were incubated at 30°C for 3 d before being imaged. WT, wild-type. (C) Fingerprint plots of the 25 thioredoxin-domain containing genes screened. The boxes are colored according to whether mutations in these genes leads to sensitization (orange) or resistance (blue) in any of the 28 CRISPR screens. (D) Immunoblotting to confirm successful knock-out of *TXNDC17* in two independent clones of RPE1 hTERT *p53^-/-^* Cas9 cells. Tubulin was used as a loading control. (E) Quantitation of clonogenic survival of RPE1 cells of the indicated genotypes in response to increasing doses of MNNG, MMS, cisplatin and KBrO_3_. (F) Competitive growth assays in untreated RPE1 cells transduced with virus expressing the indicated sgRNAs. Data represent mean fraction of GFP-positive cells ± SD normalized to day 0 (*N*= 3, independent transductions). TIDE analysis is shown in Table S6.

**Figure S7.**
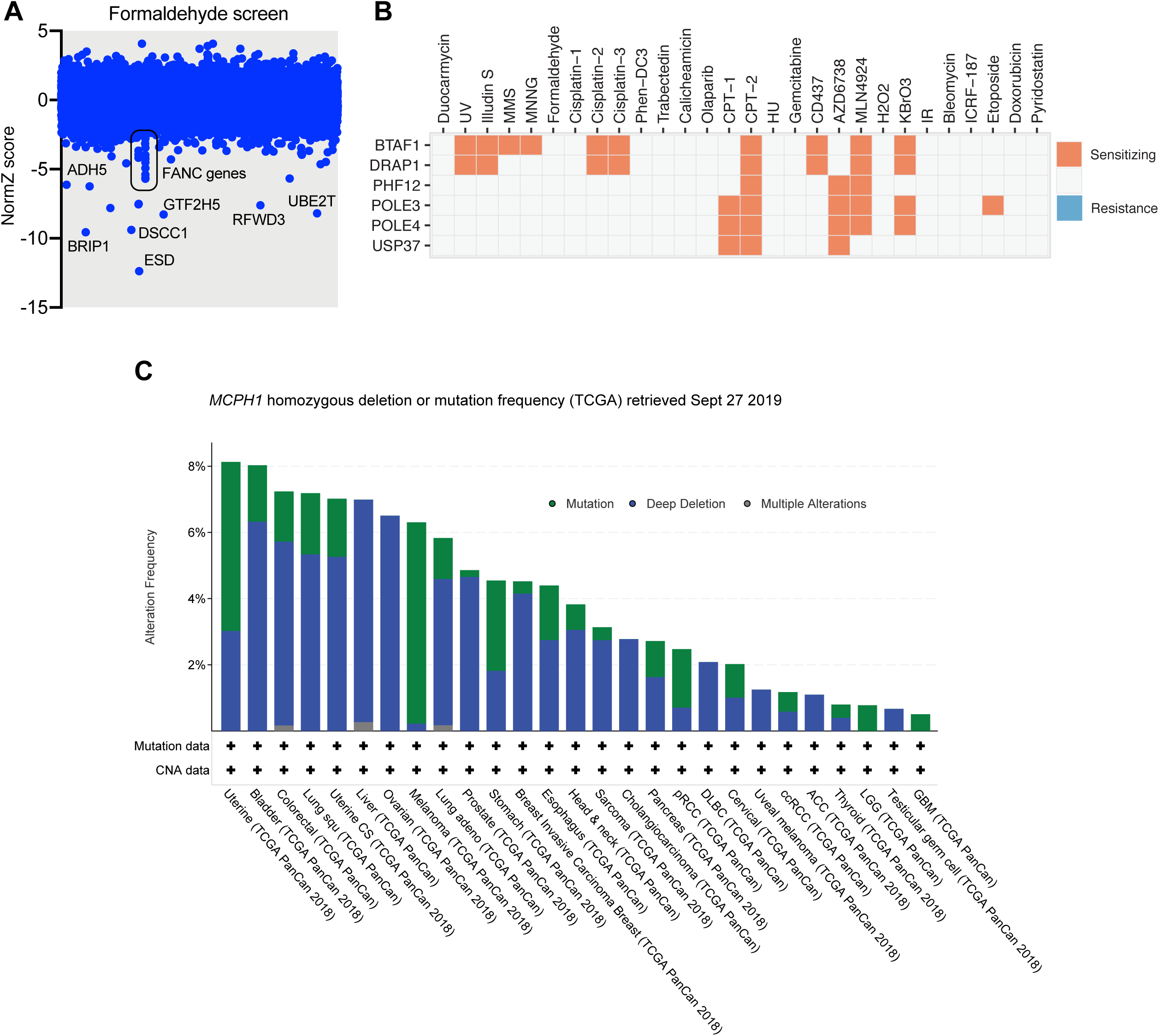
ESD, BTAF1, DRAP1 and MCPH1 as genes of interest. (A) CRISPR dropout screen results for RPE1 cells exposed to formaldehyde. Some of the top hits involved in the formaldehyde detoxification and the formaldehyde-induced DNA lesions repair have been highlighted. (B) Fingerprint plots of *BTAF1*, *DRAP1*, *PHF12*, *POLE3*, *POLE4* and *USP37*. The boxes are colored according to whether mutations in these genes leads to sensitization (orange) or resistance (blue) in any of the 28 CRISPR screens. (C) Histogram plot generated from data at cBioPortal (http://www.cbioportal.org) showing the frequency of *MCPH1* deletions and mutations across different types of cancer.

